# Sugar transporters enable a leaf beetle to accumulate plant defense compounds

**DOI:** 10.1101/2021.03.03.433712

**Authors:** Zhi-Ling Yang, Hussam Hassan Nour-Eldin, Sabine Hänniger, Michael Reichelt, Christoph Crocoll, Fabian Seitz, Heiko Vogel, Franziska Beran

## Abstract

Many herbivorous insects selectively accumulate plant toxins for defense against predators; however, little is known about the transport processes that enable insects to absorb and store defense compounds in the body. Here, we investigate how a specialist herbivore, the horseradish flea beetle, accumulates high amounts of glucosinolate defense compounds in the hemolymph. Using phylogenetic analyses of coleopteran membrane transporters of the major facilitator superfamily, we identified a clade of glucosinolate-specific transporters (*Pa*GTRs) belonging to the sugar porter family. *PaGTR* expression was predominantly detected in the excretory system, the Malpighian tubules. Silencing of *PaGTR*s led to elevated glucosinolate excretion, significantly reducing the levels of sequestered glucosinolates in beetles. This suggests that *Pa*GTRs reabsorb glucosinolates from the Malpighian tubule lumen to prevent their loss by excretion. Ramsay assays performed with dissected Malpighian tubules confirmed a selective retention of glucosinolates. Thus, the selective accumulation of plant defense compounds in herbivorous insects can depend on the ability to prevent excretion.

## Introduction

Physiological and chemosensory adaptations of herbivorous insects to plant defense compounds have played an important role in the evolution of insect-plant interactions and insect host range^1–4^. Among the most remarkable insect adaptations is the ability to selectively accumulate (sequester) plant defense compounds for protection from generalist predators^5–7^. Indeed, it is increasingly recognized that predator pressure has promoted insect adaptations to plant defenses and driven the evolution of specialized host plant associations^7–10^. Sequestration evolved in all major clades of herbivorous insects and is particularly wide-spread in the megadiverse phytophagous Coleoptera and Lepidoptera^11^; however, the mechanisms by which sequestering insects can accumulate ingested plant defense compounds in their body remain poorly understood.

Glucosinolates are hydrophilic defense compounds produced by plants of the order Brassicales that, together with the plant β-thioglucosidase enzyme myrosinase, constitute an activated two component defense system^12–14^. When tissue damage brings both components together, glucosinolates are rapidly hydrolyzed to unstable aglucones that give rise to different products^15^ of which isothiocyanates are the most detrimental to small herbivores^16–19^. Some insects can prevent glucosinolate hydrolysis and accumulate ingested glucosinolates in the body^20–25^. A defensive function of glucosinolate sequestration has been demonstrated in specialist aphids and flea beetles, which convergently evolved insect myrosinases enabling glucosinolate activation in response to predator attack^21,23,26^. The turnip sawfly, *Athalia rosae*, sequesters glucosinolates but does not possess own myrosinase activity, making a role in defense less likely^27,28^. Instead, a rapid absorption of ingested glucosinolates across the gut epithelium could represent a detoxification mechanism as it spatially separates glucosinolates from the co-ingested plant myrosinase in the gut lumen^28,29^. Due to their physicochemical properties, transport of glucosinolates and other plant glucosides such as cyanogenic and phenolic glucosides is proposed to be mediated by membrane carriers^6,23,29^. For example, Strauss and coworkers demonstrated the role of an ATP-binding cassette (ABC) transporter in the sequestration of the phenolic glucoside salicin in the defense glands of the poplar leaf beetle, *Chrysomela populi*^30^. More recently, Kowalski and coworkers identified an ABC transporter in the dogbane leaf beetle, *Chrysochus auratus*, with high activity towards a plant cardenolide, which adult beetles sequester in defense glands^31^. Apart from these examples, no other membrane transporters involved in sequestration have been identified to date.

We previously investigated the sequestration of glucosinolates in the flea beetle *Phyllotreta armoraciae*^25,26^. This species is monophagous on horseradish in nature, but feeds on various brassicaceous plants in the laboratory^32–34^ and sequesters glucosinolates mainly in the hemolymph^25^. *P. armoraciae* prefer to sequester aliphatic glucosinolates over indolic glucosinolates; however, the sequestration pattern depends both on the levels and composition of glucosinolates in the food plant. Interestingly, the beetles also selectively excrete sequestered glucosinolates to balance the accumulation of new glucosinolates and maintain stable glucosinolate levels around 35 nmol per mg beetle fresh weight^25^. Based on these findings, we hypothesize that glucosinolate transporters are localized in gut and Malpighian tubule epithelial membranes in *P. armoraciae*.

Here we use a comparative phylogenetic approach to identify candidate glucosinolate transporter genes within the major facilitator superfamily. We focus on a *P. armoraciae*-specific clade comprising 21 putative sugar porters, which are predominantly expressed in the Malpighian tubules, and demonstrate glucosinolate-specific import activity for 13 transporters. We investigate the role of the identified glucosinolate transporters *in vivo* using RNA interference and establish the transport mechanism of one transporter selected as a model *in vitro*. Our results suggest that transporter-mediated reabsorption of glucosinolates in the Malpighian tubules enables *P. armoraciae* to sequester high amounts of glucosinolates in the hemolymph.

## Results

### Selection of candidate transporters

To elucidate the molecular basis of glucosinolate transport in *P. armoraciae*, we generated a gut- and Malpighian tubule-specific transcriptome. In this transcriptome, we predicted a total of 1401 putative membrane transporters using the transporter automatic annotation pipeline (TransAAP)^35^. Of these, 353 are putative members of the major facilitator superfamily (MFS; Transporter Classification (TC)# 2.A.1; Supplementary Table 1), which are known to transport a broad spectrum of substrates including sugars and amino acids across membranes^36,37^.

We selected MFS transporters as our candidates and additionally assumed that gene duplications have played a role in the evolution of glucosinolate transport activity in *P. armoraciae*.To detect species-specific expansions of MFS transporters, we annotated MFS transporters in the publicly available genomes of three non-sequestering beetle species (Supplementary Data 1). By phylogenetic analysis, we identified the largest *P. armoraciae*-specific clade comprising 21 putative sugar porters (TC# 2.A.1.1) for further study (Supplementary Data 2). Members of this clade share between 31 and 93% amino acid sequence identity and are most similar to insect trehalose transporters.

### Functional characterization of candidate transporters

We expressed the candidate transporters in High Five insect cells and screened them for glucoside import activity. In addition to glucosinolates, we tested different types of non-host glucosides as substrates, namely iridoid, cyanogenic, and phenolic glucosides (Fig. 1a). Insect cells expressing 15 different transporters accumulated significantly higher levels of at least one tested glucoside compared to control cells. Of these, 13 transporters were specific for glucosinolates (Fig. 1b, c), hereafter referred to as *Phyllotreta armoraciae* Glucosinolate Transporter (*Pa*GTR) 1 to 13. Most *Pa*GTRs showed broad and overlapping substrate spectra; for example, 10 different *Pa*GTRs used 2-propenyl glucosinolate, the major glucosinolate in the host plant of *P. armoraciae*, as a substrate. The other two transporters mediated iridoid glucoside uptake into insect cells, whereas phenolic and cyanogenic glucosides were not used as substrates by any of the recombinant transporters (Fig. 1c and Supplementary Fig. 1).

**Fig. 1.**
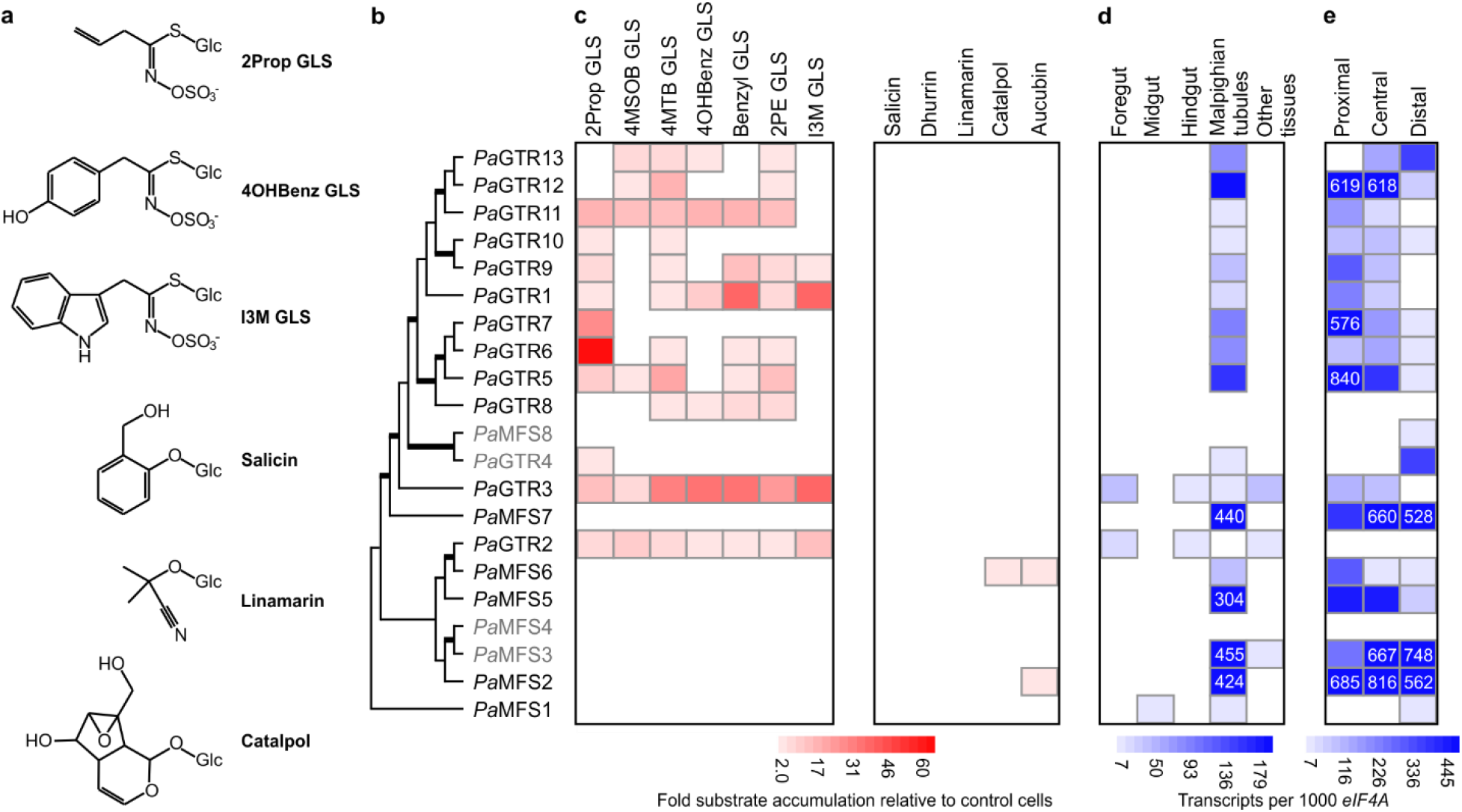
Import activity and expression pattern of candidate MFS transporters. **a** Chemical structures of selected glucosinolates (GLS) and non-host glucosides used in transport activity assays. **b** Phylogenetic relationships of candidate transporters selected based on the diversification pattern of coleopteran MFS transporters shown in Supplementary Data 2. Branches shown in bold have a bootstrap support higher than 95%. Recombinant proteins expressed in High Five insect cells were detected by Western blotting. For four candidates, we did not detect recombinant protein (Supplementary Fig. 1); the names of these candidates are written in grey **c** Recombinant transporters were screened for glucoside uptake activity in assays using equimolar mixtures of glucosinolates or non-host glucosides. Glucoside accumulation in transfected insect cells is expressed relative to that in mock-transfected insect cells used as background control. Values represent the mean of three assays. **d** Expression pattern of candidate MFS transporter genes in different tissues of *P. armoraciae*. **e** Expression pattern of candidate MFS transporter genes in different regions of the Malpighian tubule. Copy number estimates are given per 1000 copies of mRNA of the reference gene *eIF4A*. Low gene expression levels are visualized by limiting the scale to a value of 200 and 500 in **d** and **e**, respectively. The exact values are provided for genes with higher expression level. Each value represents the mean of four and three biological replicates in **d** and **e**, respectively. 2Prop, 2-propenyl; 4MSOB, 4-methylsulfinylbutyl; 4MTB, 4-methylthiobutyl; 4OHBenz, 4-hydroxybenzyl; 2PE, 2-phenylethyl; I3M, indol-3-ylmethyl.

### Tissue-specific expression of *PaGTR*s

To determine the localization of *Pa*GTRs in *P. armoraciae*, we compared the transcript levels in foregut, midgut, hindgut, Malpighian tubules, and other tissues by quantitative PCR. All except two *PaGTR*s were specifically expressed in the Malpighian tubules (Fig. 1d and Supplementary Fig. 2). Along the tubule, *PaGTR*s were predominantly expressed in the proximal region close to the midgut-hindgut junction (Fig. 1e and Supplementary Fig. 2d). This Malpighian tubule region was previously shown to reabsorb metabolites to prevent their loss by excretion^38–40^.

### Function of *Pa*GTRs *in vivo*

The Malpighian tubules are bathed in hemolymph, the major storage site for sequestered glucosinolates in *P. armoraciae*. We analyzed the glucosinolate levels in the hemolymph and detected total concentrations of up to 106 mM (Supplementary Table 2). Which role does *Pa*GTR-mediated glucosinolate transport in Malpighian tubules play in glucosinolate storage? We addressed this question by silencing *PaGTR* expression in adult *P. armoraciae* beetles. We selected *Pa*GTR1 as a model because this transporter prefers indol-3-ylmethyl (I3M) glucosinolate, a substrate that is transported by only one other Malpighian tubule-specific transporter (Fig. 1c). After confirming the specific silencing of *PaGTR1* expression by quantitative PCR (Fig. 2a; Supplementary Fig. 3a), we fed the beetles with leaves of *Arabidopsis thaliana* Col-0 (*Arabidopsis*) containing I3M glucosinolate.

**Fig. 2.**
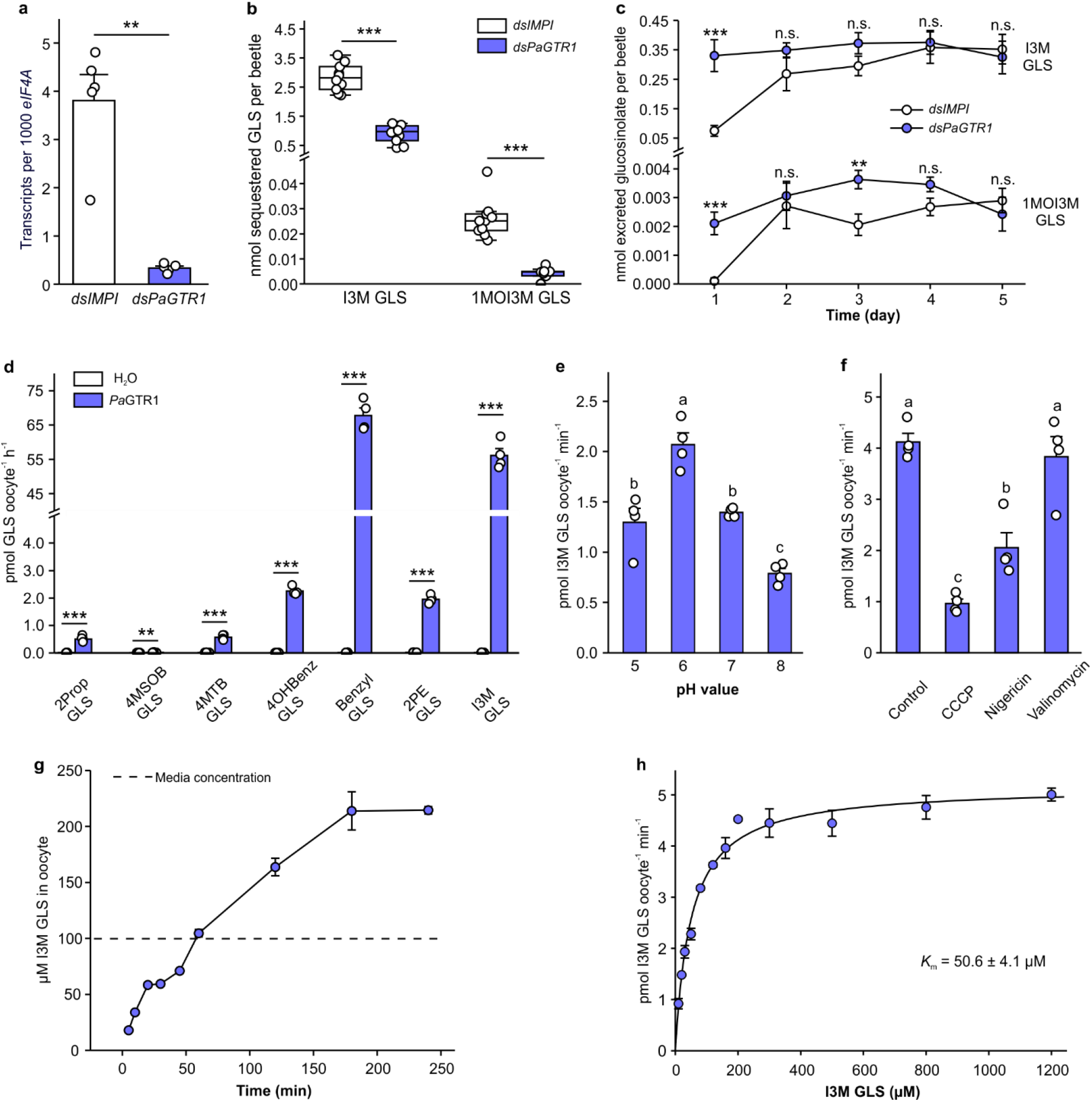
Functional and biochemical characterization of *Pa*GTR1-mediated glucosinolate transport. **a** *PaGTR1* expression in *P. armoraciae* four days after dsRNA injection (*n* = 5). **b** Accumulation of indol-3-ylmethyl (I3M) GLS and 1-methoxyindol-3-ylmethyl (1MOI3M) GLS in adult *P. armoraciae* beetles after five days feeding on *Arabidopsis* leaves. Box plots show the median, interquartile range, and outliers of each data set (*n* = 10). **c** Excreted amounts of I3M GLS and 1MOI3M GLS over five days feeding on *Arabidopsis* (*n* = 9 for day 4, *n* =10 for other days). **d** Substrate preference of *Pa*GTR1. Oocytes expressing *Pa*GTR1 and control oocytes were incubated with equimolar mixtures of seven glucosinolates (*n* = 4). **e** pH dependency of *Pa*GTR1 (*n* = 4). **f** Effects of different ionophores on *Pa*GTR1-mediated glucosinolate transport (*n* = 4). **g** Time course of glucosinolate accumulation in *Xenopus* oocytes expressing *Pa*GTR1 (*n* =4). **h** Kinetic analysis of *Pa*GTR1-mediated I3M GLS transport (*n* = 4). Data are shown as mean ± s.e.m. Treatments were compared by two-tailed Student’s *t*-test, Mann-Whitney *U* test or the method of generalized least squares (**a**, **b**, **c**, **d**), or one-way analysis of variance (ANOVA) (**e**, **f**). Bars labeled with different letters are significantly different (*P* < 0.05). n.s., not significantly different, ** *P* < 0.01, *** *P* < 0.001.

Although the treatment did not influence beetle feeding behavior (Supplementary Fig. 3b), we detected significantly less I3M glucosinolate and 1-methoxyindol-3-ylmethyl (1MOI3M) glucosinolate in *PaGTR1*-silenced beetles than in control beetles (Fig. 2b). In addition, *PaGTR1*-silencing led to elevated levels of I3M and 1MOI3M glucosinolates in the feces (Fig. 2c). The levels of other glucosinolates were not reduced in *PaGTR1*-silenced beetles compared to the control (Supplementary Table 3 and Supplementary Fig. 4). The effect of *PaGTR1* knock-down on the accumulation and excretion of specific glucosinolates in *P. armoraciae* is consistent with the substrate preference of recombinant *Pa*GTR1 in insect cell-based uptake assays, and suggests that *Pa*GTR1 prevents excretion of two indolic glucosinolates by reabsorbing them from the Malpighian tubule lumen.

### *Pa*GTR1 is a proton-dependent high affinity glucosinolate transporter

To establish the mechanism of *Pa*GTR1-mediated glucosinolate reabsorption, we expressed *PaGTR1* in *Xenopus laevis* oocytes and analyzed its biochemical properties. First, we analyzed the substrate preference of oocyte-expressed *Pa*GTR1, and found it to be similar to that of insect cell-expressed *Pa*GTR1 (Fig. 2d). Previously characterized members of the sugar porter family act either as uniporters or as proton-dependent active transporters^41–43^; thus, we analyzed whether the extracellular pH influences I3M glucosinolate accumulation in *Pa*GTR1-expressing oocytes. The glucosinolate uptake rate depended on the external pH and was maximal at pH 6, which is consistent with the acidic pH in the Malpighian tubule lumen of *P. armoraciae* (Fig. 2e and Supplementary Fig. 5). Uptake activity decreased significantly when pH was greater than 6, suggesting that *Pa*GTR1-mediated glucosinolate transport is driven by a proton gradient across the membrane. To test this, we performed glucosinolate uptake assays at pH 6 and added either the protonophore carbonyl cyanide *m*-chlorophenyl hydrazine (CCCP), the K^+^/H^+^ exchanger nigericin, or the K^+^ ionophore valinomycin to the assay medium. Addition of CCCP and nigericin decreased the glucosinolate uptake rates by 77% and 50% compared to control assays, respectively, whereas valinomycin did not influence transport by *Pa*GTR1-expressing oocytes. The effects of different ionophores on glucosinolate transport show that *Pa*GTR1-mediated glucosinolate uptake is proton-driven, but is not influenced by K^+^ (Fig. 2f).

A time-course analysis of I3M glucosinolate accumulation in *Pa*GTR1-expressing oocytes revealed that glucosinolate levels saturated after three hours at an intracellular concentration that was twofold higher than in the assay buffer. This result confirmed that *Pa*GTR1 can mediate glucosinolate uptake against a concentration gradient (Fig. 2g). Plotting the transport rate as a function of increasing glucosinolate concentration yielded a saturation curve that was fitted to Michaelis-Menten kinetics with a *K*_m_ value of 50.6 ± 4.1 μM (Fig. 2h). Considering the much higher I3M glucosinolate concentration of 2082 ± 668 μM in the hemolymph (Supplementary Table 2), our results characterize *Pa*GTR1 as a high affinity proton-dependent glucosinolate transporter in *P. armoraciae*.

### *P. armoraciae* selectively retain glucosinolates

We further investigated the role of selective transport in Malpighian tubules by analyzing the excretion of two glucosinolates (2-propenyl- and 4-hydroxybenzyl glucosinolate) and three non-host glucosides (salicin, linamarin and catalpol) after injection into the hemolymph of adult *P. armoraciae* beetles. Within one day, beetles excreted between 18% and 58% of the injected non-host glucosides and less than 2% of the injected glucosinolates (Fig. 3a). In addition, we compared the recovery of injected glucosides after 30 mins with that after one day. While the levels of glucosinolates remained stable, we recovered significantly less non-host glucosides from beetle bodies. However, significantly more catalpol than salicin and linamarin was recovered (Fig. 3a), suggesting that catalpol was excreted or metabolized at a lower rate than the other two non-host glucosides.

**Fig. 3.**
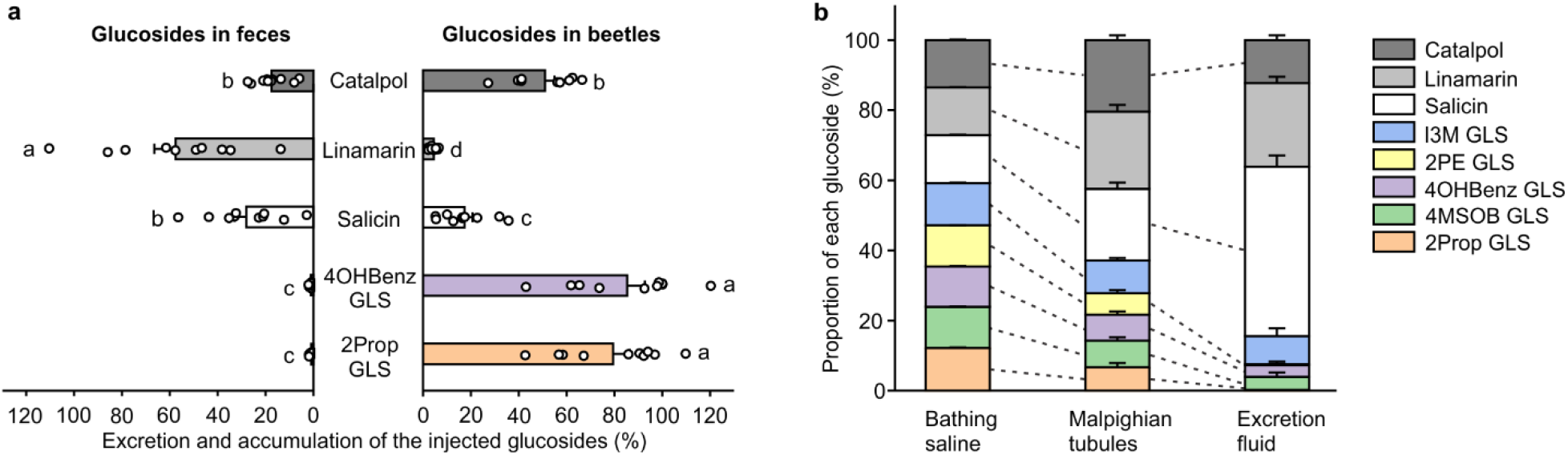
Selective excretion of plant glucosides *in vivo* and *in situ*. **a** Excretion and accumulation of plant glucosides injected in beetles. Each beetle was injected with 100 nL of an equimolar mixture of five glucosides. Injected beetles were sampled 30 min after the injection and after one day feeding on *Arabidopsis (n* = 10). Glucoside excretion was analyzed by quantifying the amount of excreted glucosides in the feces (*n* = 10). Glucoside content in beetles and feces is expressed relative to the glucoside amounts detected in beetles 30 min after injection (set to 100%). The relative accumulation and excretion of glucosides was compared using the method of generalized least squares, respectively. Bars labeled with different letters are significantly different (*P* < 0.05). **b** Excretion of plant glucosides by isolated Malpighian tubules. The dissected Malpighian tubule was placed in a droplet of saline and a mixture of eight different plant glucosides each at a concentration of 6.7 mM. To visualize excretion, we added 0.1% (w/v) amaranth. After 2-3 h, bathing saline, Malpighian tubule and excretion fluid were sampled, extracted and analyzed by LC-MS/MS. Paired *t*-tests were used to compare the relative composition of plant glucosides in bathing saline and Malpighian tubule, and Malpighian tubule and excretion fluid, respectively. Dashed lines indicate significant differences between samples (*P* < 0.05, *n* = 11). Data are shown as mean ± s.e.m.

To confirm that the selective retention of glucosinolates in *P. armoraciae* is due to reabsorption, we analyzed glucoside excretion *in situ* using Ramsay assays (Supplementary Fig. 6). We exposed dissected Malpighian tubules to an equimolar mixture of glucosides and compared the glucoside composition in the bathing saline and Malpighian tubule tissue, and in the Malpighian tubule tissue and the excretion fluid, respectively. The Malpighian tubule tissue contained significantly lower proportions of glucosinolates than the bathing saline and correspondingly higher proportions of non-host glucosides (Fig. 3b). The excretion fluid contained even lower percentages of most glucosinolates, in particular of 2-propenyl and 2-phenylethyl glucosinolates. In addition, we detected less catalpol in the excretion fluid than in the Malpighian tubule tissue. Combined, these findings show that the Malpighian tubules of *P. armoraciae* selectively reabsorb glucosinolates and catalpol, which is consistent with the substrate spectra of recombinant candidate transporters (Fig. 1c).

### *Pa*GTRs prevent excretion of 2-propenyl glucosinolate

2-Propenyl glucosinolate represents the major glucosinolate in the host plant of *P. armoraciae* and is sequestered at very high levels in the beetle hemolymph (Supplementary Table 2). To determine whether the identified *Pa*GTRs also capable of reabsorbing 2-propenyl glucosinolate in *P. armoraciae*, we simultaneously silenced the expression of a clade comprising four Malpighian tubule-specific transporters (*PaGTR5/6/7/8*),among which three used 2-propenyl glucosinolate as a substrate *in vitro* (Fig. 1b, c). Beetles with silenced *PaGTR5/6/7/8* expression excreted significantly more 2-propenyl glucosinolate than control beetles (Fig. 4a, c and Supplementary Fig. 7, 8). In addition, 2-propenyl glucosinolate levels were significantly lower in *PaGTR5/6/7/8*-silenced beetles than in control beetles (Fig. 4b and Supplementary Table 4). These results support our hypothesis that the major function of Malpighian tubule-expressed *Pa*GTRs is reabsorption.

**Fig. 4.**
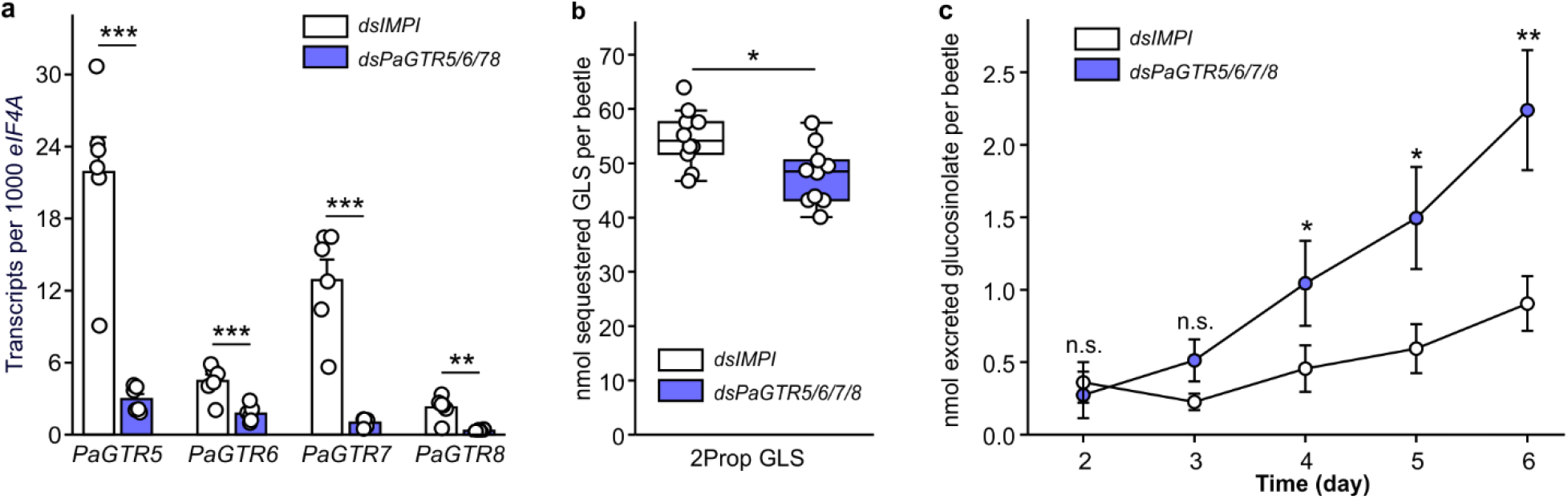
In vivo functional characterization of glucosinolate transport mediated by *PaGTR5/6/7/8*. **a** *PaGTR5/6/7/8* expression in *P. armoraciae* six days after dsRNA injection (*n* = 6). **b** Accumulation of 2-propenyl glucosinolate (2Prop GLS) in adult *P. armoraciae* beetles after six days feeding on *Arabidopsis* leaves followed by one day starvation. Box plots show the median, interquartile range, and outliers of each data set (*n* = 10). **c** Excreted amounts of 2Prop GLS from day 2 until day 6 of feeding on *Arabidopsis* (*n* = 10). n.s., not significantly different; \**P* < 0.05, \*\**P* < 0.01, \*\*\**P* < 0.001. Data are shown as mean ± s.e.m.

### Silencing of *PaGTR2* and *PaGTR3* does not affect glucosinolate accumulation from *Arabidopsis*

Two transporter genes, *PaGTR2* and *PaGTR3*, were found to be expressed in the foregut, hindgut, and other tissues. Because of their expression in the foregut, we investigated whether *Pa*GTR2 and *Pa*GTR3, which showed a broad substrate specificity *in vitro*, play a role in glucosinolate absorption from the gut lumen. However, silencing of *PaGTR2* and *PaGTR3* expression did not influence the accumulation of ingested glucosinolates in beetles in feeding assays with *Arabidopsis* (Supplementary Fig. 9).

## Discussion

In this study, we identified glucosinolate-specific transporters in *P. armoraciae* that play an important role in sequestration. We show that *PaGTR*s are mainly expressed in the Malpighian tubules and mediate glucosinolate import *in vitro*. Silencing *PaGTR* expression in Malpighian tubules led to increased glucosinolate excretion, which indicates that *Pa*GTRs reabsorb glucosinolates from the Malpighian tubule lumen into the epithelial cells. These findings demonstrate that active reabsorption in the Malpighian tubules is required for glucosinolate accumulation in the hemolymph of *P. armoraciae*.

Glucosinolate transporters have previously been identified in the model plant *Arabidopsis* by screening 239 transport proteins for glucosinolate uptake activity in *Xenopus* oocytes^44^. Plant GTRs belong to the nitrate/peptide family (NPF) and presumably evolved from ancestral cyanogenic glucoside transporters^45^. The corresponding transporter family in *P. armoraciae*, annotated as proton-dependent oligopeptide transporter family (POT), comprised only six putative transporters (Supplementary Table 1) and was not investigated in our study. Instead, our phylogenetic analysis of coleopteran MFS transporters provided evidence for species-specific expansions in particular within the sugar porter family, which led us to focus on the largest *P. armoraciae*-specific clade of putative sugar porters. All *Pa*GTRs except for *Pa*GTR2 clustered together in one well-supported group (Fig. 1b). Separate of this group we additionally identified two iridoid glycoside-specific transporters. Our study thus highlights the potential of sugar porters to evolve activity towards glycosylated defense compounds.

The presence of iridoid glycoside-specific transporters in *P. armoraciae* was surprising, as plants of the order Brassicales do not produce iridoid glycosides^46,47^. Consistently, we observed reabsorption of the iridoid glycoside catalpol in Malpighian tubules (Fig. 3). Several flea beetles of the genus *Longitarsus* selectively sequester aucubin and catalpol^48^, but this genus is likely not closely related to *Phyllotreta*^49^. Nevertheless, it is possible that iridoid glycoside-specific transporters that had evolved in an ancestor of *Phyllotreta* have retained their function even after flea beetles have adapted to plant families with distinct defense compounds.

An evolutionary link between the sequestration of iridoid glucosides and glucosinolates has previously been discovered in the sawfly genus *Athalia* in which a host shift occurred from iridoid glycoside-containing Lamiales to glucosinolate-containing Brassicaceae^50^. Sequestration experiments demonstrated that Brassicaceae-feeders were able to sequester both iridoid glycosides and glucosinolates, whereas Lamiales-feeders only sequestered iridoid glycosides. Based on these findings, Opitz and coworkers hypothesized that glycoside transporters evolved a broader substrate specificity in Brassicaceae feeders. It is also imaginable that the presence of iridoid glycoside-specific transporters in the ancestor of *Phyllotreta* has facilitated the evolution of glucosinolate transport activity when flea beetles specialized on brassicaceous plants. To better understand the evolutionary origin of *Pa*GTRs, it will be interesting to investigate the evolution and function of sugar porters across different flea beetle genera.

The specific expression of most *PaGTR*s in the Malpighian tubules indicated an important role of the beetle’s excretory system in sequestration. In fact, pioneering work by Meredith and coworkers published in 1984 demonstrated that the Malpighian tubules of the large milkweed bug, *Oncopeltus fasciatus*, actively reabsorb the polar cardiac glycoside ouabain, which is sequestered predominantly in the integument, after its passive secretion into the Malpighian tubule lumen^38^. Although it has generally been recognized that sequestering insects require mechanisms to prevent the excretion of polar metabolites from the hemolymph^51^, these mechanisms have not been further investigated. Our physiological assays demonstrate that the Malpighian tubule epithelium of *P. armoraciae* discriminates among different types of glucosinolates and non-host glucosides and can reabsorb in particular glucosinolates against a concentration gradient (Fig. 3). Since *Pa*GTRs mediated glucosinolate import *in vitro*, we propose that they are located at the apical membrane and thereby enable glucosinolate uptake from the lumen. To complete transepithelial glucosinolate transport, a so far unknown membrane transporter localized at the basolateral membrane is required to export glucosinolates into the hemolymph (Fig. 5).

**Fig. 5.**
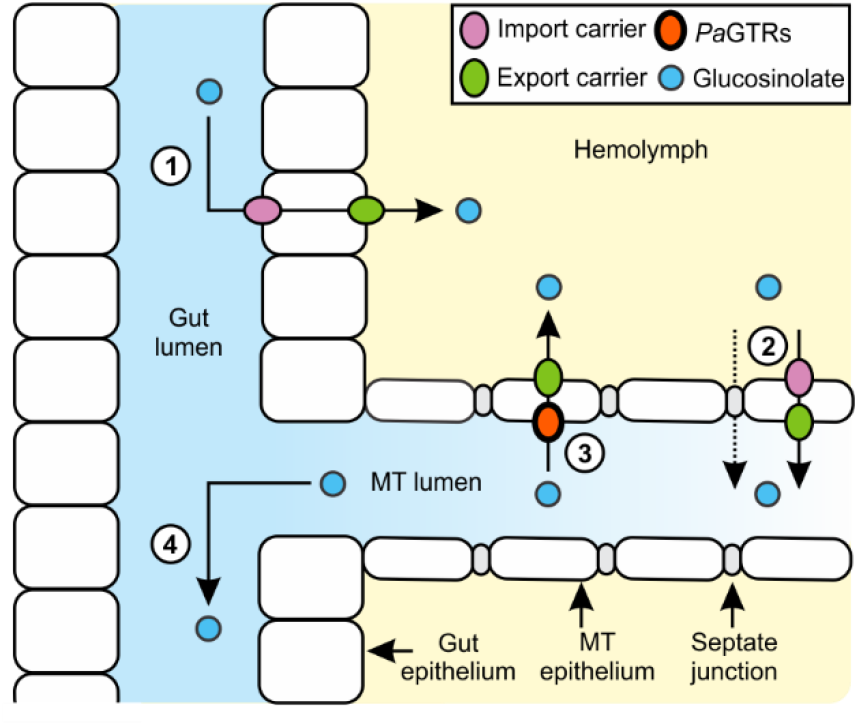
Proposed model of glucosinolate sequestration in adult *P. armoraciae*. (1) Absorption of ingested glucosinolates from the gut lumen into the hemolymph; (2) Transcellular or paracellular glucosinolate transport from the hemolymph into the Malpighian tubule (MT) lumen; (3) Selective reabsorption of glucosinolates from the MT lumen into the hemolymph; (4) Excretion of excess glucosinolates. *Pa*GTRs localized in the MT are proposed to import glucosinolates from the lumen across the apical membrane into the epithelial cells.

The expression profile of putative sugar porter genes in *P. armoraciae* (Supplementary Data 1) and a recent proteomic study performed with the poplar leaf beetle *C. populi*^52^ showed that many sugar porters are specifically expressed in the Malpighian tubules. To our knowledge, only one Malpighian tubule-specific sugar porter has been characterized to date - sugar transporter 8 from the brown plant hopper, *Nilaparvata lugens*, which reabsorbs trehalose^42^. Thus, there is first evidence that sugar porters localized in Malpighian tubules play a role in reabsorption. The localization and function of putative sugar porters we have identified in published beetle genomes has not yet been investigated; however, a phylogenetic analysis suggests that the Malpighian tubule-specific sugar porter CpSP-like17 from *C. populi* and *Pa*GTRs evolved from a common ancestor (Supplementary Fig. 10). We thus propose that *Pa*GTRs evolved by duplications of a Malpighian tubule-localized sugar porter. Investigating the localization of flea beetle sugar porters that are phylogenetically closely related to *Pa*GTRs will help to establish whether *Pa*GTRs indeed evolved from Malpighian tubule-specific sugar porters.

The distinct expression pattern of *PaGTR2 and PaGTR3* (Fig. 1d) suggested that *Pa*GTRs are not only involved in reabsorption in the Malpighian tubules. Here, the expression in the foregut suggested for example a role in glucosinolate absorption. Although the insect midgut has been proposed to be the major tissue involved in absorption^40^, there is initial evidence that glucosinolate absorption might already occur in the foregut of *A. rosae* larvae^29^. However, as silencing *PaGTR2* and *PaGTR3* expression had no effect on the accumulation of ingested glucosinolates in beetles, it is likely that additional or other membrane transporters are required. To establish the mechanism of glucosinolate absorption, we currently investigate the localization and specificity of glucosinolate uptake in adult *P. armoraciae*.

Transport processes play a central role in the evolution of insect sequestration mechanisms. Our study highlights that sequestering insects require mechanisms that prevent the excretion of target compounds, for example by active reabsorption. We furthermore show that in *P. armoraciae*, reabsorption is selective and thus can control not only the level but also the composition of sequestered metabolites in the beetle. Understanding the molecular basis of sequestration now opens up the possibility to manipulate the accumulation of defense compounds in insects in order to investigate how sequestration influences trophic interactions and shapes the composition of ecological communities.

## Methods

### Insect culture

*P. armoraciae* beetles were reared on potted *Brassica juncea* cultivar “Bau Sin” plants or on potted *Brassica rapa* cultivar “Yo Tsai Sum” plants (Known-You Seed Co. Ltd.) in mesh cages (Bugdorm, MegaView Science Co., Ltd.) in a controlled environment chamber at 24 °C, 60% relative humidity and a 16-h photoperiod. Food plants were cultivated in a growth chamber at 24 °C, 55% relative humidity, and a 14-h photoperiod. Beetles were provided with three-to four-week old plants once per week, and plants with eggs were kept separately for larval development. Larvae were allowed to pupate in the soil and after three weeks, the soil with pupae was transferred into plastic boxes (9 L, Lock&Lock) until the new generation of beetles emerged.

### RNA isolation, RNAseq and *de novo* transcriptome assembly

Total RNA was extracted from dissected foregut, midgut, hindgut, and Malpighian tubule tissue of newly emerged *P. armoraciae* beetles that were reared on *B. juncea* using the innuPREP RNA Mini Kit (Analytik Jena). Tissues from at least 10 beetles were pooled per sample. RNA integrity was verified using an Agilent Technologies 2100 Bioanalyzer with the RNA 6000 Nano Kit (Agilent Technologies). RNA quantity was determined using a Nanodrop ND-1000 spectrophotometer (PEQlab Biotechnologie GmbH). One set of RNA samples was sequenced by GATC Biotech on the HiSeq 2500 System from Illumina in Rapid Run mode, using the paired-end (2 × 125 bp) read technology at a depth of 15-25 million reads for each sample. For a second set of samples consisting of four biological replicates per tissue, we additionally performed an on-column DNA digestion with the innuPREP DNase I Digest Kit (Analytik Jena) according to the manufacturer’s instructions. RNA samples were poly(A)-enriched, fragmented, and sequenced at the Max Planck Genome Centre Cologne on the HiSeq 3000 Sequencing System from Illumina, using the paired-end (2 × 150 bp) read technology at a depth of 22 million reads for each sample. Sequencing reads were filtered to remove bad-quality reads based on fastq file scores and trimmed based on read length using CLC Genomics Workbench software version 10.1. With a randomly sampled set of 420 million reads from the two sets of sequencing data, a transcriptome was assembled *de novo* with the following parameters: nucleotide mismatch cost = 1; insertion = deletion costs = 2; length fraction = 0.6; similarity = 0.9. Conflicts among the individual bases were resolved by voting for the base with the highest frequency. After removing contigs shorter than 250 bp, the final assembly contained 36,445 contigs with an N50 contig size of 2,115 bp.

### Identification of coleopteran MFS transporters

We predicted a protein dataset for *P. armoraciae* by translating each contig of the gut and Malpighian tubule-specific transcriptome into all six reading frames. After removing sequences shorter than 50 amino acids, we submitted the protein dataset (267,568 sequences) to the Transporter Automatic Annotation Pipeline (TransAAP) hosted at the TransportDB 2.0 web portal^35^. This initial annotation predicted a total of 1,401 putative transporter sequences and revealed the major facilitator superfamily (MFS) and the ATP-binding cassette (ABC) transporters to be the largest transporter families (Supplementary Table 1). We focused on MFS transporters, which are classified into more than 80 families^53^. We used one protein sequence from each MFS family as query to search candidate MFS transporters in the protein dataset from *P. armoraciae* using Blastp (E-value threshold of 10^−5^), and assigned each candidate to an MFS family based on sequence similarity to transporter sequences deposited in TCDB. Additional candidates were identified by repeating the search procedure with an extended dataset including the candidate MFS transporters from *P. armoraciae*. The number of TMDs for each candidate was predicted using the TMHMM Server v.2.0^54^. Partial sequences encoding less than six predicted TMDs were removed from the dataset. The same strategy was used to identify putative MFS transporters in protein datasets that were predicted from the genomes of *Leptinotarsa decemlineata*, genome annotations v0.5.3^55^, *Anoplophora glabripennis*, assembly Agla_1.0^56^, and *Tribolium castaneum*, assembly Tcas3.0^57^, respectively. The predicted protein sequences are provided in Supplementary Data 1.

### Digital gene expression analysis

Digital gene expression analysis of putative MFS transporters identified in the *P. armoraciae* transcriptome was carried out using CLC Genomics workbench v 10.1 by mapping the Illumina reads from the second set of samples onto the reference transcriptome, and counting the reads to estimate gene expression levels. For the cloned major facilitator superfamily genes, complete open reading frames were used as reference sequences for mapping. For read alignment, we used the following parameters: nucleotide mismatch cost = 2; insertion = deletion costs = 3; length fraction = 0.6; similarity fraction = 0.9; maximum number of hits for a read = 15. Each pair of reads was counted as two. Biases in the sequence datasets and different transcript sizes were corrected using the TPM (transcripts per kilobase million) normalization method to obtain correct estimates for relative expression levels between samples.

### Phylogenetic analyses of coleopteran MFS transporters

We inferred the lineage-specific diversification patterns of putative MFS transporters from *P. armoraciae, L. decemlineata, A. glabripennis* and *T. castaneum* in phylogenetic analyses with two different datasets, one containing all identified MFS transporters (867 sequences), the other containing a subset of putative sugar porters (120 sequences) from the above four species and 35 sugar porters from *Chrysomela populi*^52^. The corresponding protein sequences were aligned using the MUSCLE algorithm^58^ implemented in MEGA 7 with default parameters. The alignments were trimmed manually and the best substitution models were determined using ProtTest 3.4.2^59^. Maximum-likelihood phylogenetic trees were constructed in IQ-TREE version 1.6.0^60^ using the VT+G+F substitution model with 1000 ultrafast bootstrap replicates for the full dataset, and the LG+G+F substitution model with 1000 bootstrap replicates for the subset of putative sugar porters.

### Identification and sequencing of candidate transporters

Based on our phylogenetic analysis, we selected the largest clade of putative MFS transporters that was specifically expanded in *P. armoraciae* for further studies. Transcriptome analysis revealed the presence of a pseudogene (*PaMFS9-ps*) that shares between 43 and 95% nucleotide sequence identity with members of the focal clade. The protein encoded by this pseudogene is predicted to possess only two transmembrane domains due to a premature stop codon caused by frame shift mutations in the coding sequence. To obtain the full-length open reading frames (ORFs) of partial transcripts, we performed rapid amplification of cDNA ends–PCR as described before^23^. All full length ORFs were cloned into the pCR™4-TOPO^®^TA (Thermo Fisher Scientific) for sequence verification.

### Tissue-specific expression of candidate transporters

We used quantitative PCR (qPCR) to analyze the expression of the candidate transporter genes in the foregut, midgut, hindgut, Malpighian tubules, and other tissues of one-day old adult *P. armoraciae* beetles (*n* = 4 biological replicates, each with two technical replicates), respectively. In addition, expression of candidate transporter genes was analyzed in the proximal, central, and distal Malpighian tubule regions (Supplementary Fig. 2d) dissected from four-day old adult *P. armoraciae* beetles (*n* = 3 biological replicates, each with two technical replicates). Primers (Supplementary Table 5) were designed using Primer3web version 4.1.0. Analyses of primer specificity and efficiency, RNA extraction, purification, cDNA synthesis, and qPCR were performed as described before^61^. Gene expression was normalized to the expression level of eukaryotic initiation factor 4A (*eIF4A*),which showed the lowest variability across tissues among four tested reference genes (Supplementary Table 6).

### Expression of candidate transporters in insect cells

For protein expression, we cloned each ORF without stop codon into the pIEx-4 expression vector (Novagen) in frame with the vector-encoded carboxy terminal 6 × His-tag and sequenced the resulting constructs. Primer sequences are listed in Supplementary Table 5. One construct of each candidate gene was used for transfection of High Five™ insect cells (Gibco) cultured in Express Five^®^SFM medium (Gibco) supplemented with 20 mM glutamine (Gibco) and 50 μg/mL gentamicin (Gibco). Confluent insect cells were diluted 1:5, dispensed in 500 μL-aliquots into 24-well plates, and incubated at 27 °C. On the next day, we transfected the cells using FuGENE HD Transfection Reagent (Promega) according to the manufacturer’s protocol. Cells treated with transfection reagent only were used as a negative control. After 48 h we harvested the cells for Western blotting and uptake assays.

### Western blotting

To confirm protein expression, transfected insect cells were washed twice with phosphate buffered saline (PBS; pH 7.4), collected by centrifugation, and resuspended in hypotonic buffer (20 mM Tris-HCl (pH 7.5), 5 mM EDTA, 1 mM DTT, 0.1% Benzonase nuclease (Merck Millipore) (v/v), and protease inhibitors (cComplete Mini, ETDA-free, Roche Diagnostics GmbH)). After incubation on ice for 10 min, the samples were frozen in liquid nitrogen, thawed, and centrifuged (16,000 × *g* for 15 min at 4 °C). The resulting cell pellet was resuspended in hypotonic buffer and used for Western blotting using HRP-conjugated anti-His antibody (1: 10,000; Novex, Life technologies).

### Glucoside uptake assays with transfected insect cells

Cells were washed with PBS (pH 5.5) by pipetting and incubated with different glucoside substrates at 200 μM in PBS (pH 5.5) for 1 h at 27 °C. Assays were performed with substrate mixtures containing seven different glucosinolates (2-propenyl glucosinolate (Roth), 4-methylsulfinylbutyl glucosinolate (Phytoplan), 4-methylthiobutyl glucosinolate (Phytoplan), 2-phenylethyl glucosinolate (Phytoplan), benzyl glucosinolate (Phytoplan), 4-hydroxybenzyl glucosinolate (isolated from *Sinapis alba* seeds as described before^62^), and indol-3-ylmethyl (I3M) glucosinolate (Phytoplan)), or five other plant glycosides (salicin (Sigma-Aldrich), linamarin (BIOZOL), dhurrin (Roth), catalpol (Sigma-Aldrich), and aucubin (Sigma-Aldrich). After incubation, cells were washed three times with ice-cold PBS (pH 5.5) by pipetting, collected in 300 μL 80% (v/v) methanol, frozen in liquid nitrogen, thawed, and centrifuged at 3,220 × g for 10 min at 4 °C. The supernatant was dried by vacuum centrifugation, dissolved in ultrapure water and analyzed by liquid chromatography coupled with tandem mass spectrometry (LC-MS/MS). All glucoside uptake assays were performed in triplicates. The amount of each substrate in transporter-expressing cells was compared with that detected in control cells. Transporters were considered active towards a substrate when the average amounts detected in transporter-expressing cells were at least twofold higher than those detected in control cells.

### Cloning of *PaGTR1* into the pNB1u vector and cRNA synthesis

We amplified the open reading frame of *PaGTR1* without stop codon by PCR using uracil-containing primers (Supplementary Table 5). The 3’ primer was designed to encode a Human influenza hemagglutinin (HA)-tag to enable the detection of recombinant protein by Western blotting if necessary. The pNB1u vector was digested overnight at 37 °C with PacI and Nt.BbvCI (New England Biolabs) to generate 8-nt overhangs. One microliter gel-purified PCR product (100 ng/μL) was combined with 1 μL gel-purified vector (50 ng/μL), 1 unit USER enzyme, 2 μL 5 x PCR reaction buffer and 5 μL H_2_0, incubated at 37 °C for 25 min followed by 25 min at room temperature. After transformation of chemically competent *E. coli* cells, colonies containing the appropriate insert were identified by Sanger sequencing. The DNA template for cRNA synthesis was amplified by PCR from the *X. laevis* expression construct using pNB1uf/r primers (Supplementary Table 5) and cRNA was synthesized using the mMESSAGE mMACHINE™ T7 Transcription Kit (Invitrogen) according to the manufacturer’s manual.

### Biochemical characterization of *Pa*GTR1 in *Xenopus* oocytes

The cRNA concentration was adjusted to 800 ng/μL with RNase-free water for oocyte injection.*X. laevis* oocytes (Ecocyte Bioscience) were injected with 50 nL containing 40 ng cRNA or with 50 nL pure water as a control using a Drummond NANOJECT II (Drummond Scientific Company) or a Nanoliter 2010 Injector (World Precision Instruments).

Injected oocytes were incubated in Kulori buffer (90 mM NaCl, 1 mM KCl, 1 mM MgCl_2_, 1 mM CaCl_2_ and 5 mM 4-(2-hydroxyethyl)-1-piperazineethanesulfonic acid (HEPES), pH 7.4) supplemented with 50μg/mL gentamicin at 16 °C for three days until assaying. All transport assays were performed at room temperature. Injected oocytes were pre-incubated for 5 min in Kulori buffer (90 mM NaCl, 1 mM KCl, 1 mM MgCl_2_, 1 mM CaCl_2_ and 5 mM 2-(N-morpholino)ethanesulfonic acid (MES), pH 6.0) before they were transferred into the same buffer containing the substrate(s). To determine the substrate preference of *Pa*GTR1, we incubated oocytes in Kulori buffer (pH 6.0) containing an equimolar mixture of 2-propenyl glucosinolate, 4-methylsulfinylbutyl glucosinolate, 4-methylthiobutyl glucosinolate, 2-phenylethyl glucosinolate, benzyl glucosinolate, 4-hydroxybenzyl glucosinolate, and I3M glucosinolate, each at 200μM, for 1 h.

The pH dependency of glucosinolate transport was determined by incubating the injected oocytes with 100 μM I3M glucosinolate for 10 min in Kulori buffer adjusted to different pH values. The effects of different ionophores on glucosinolate transport was studied by incubating the *Pa*GTR1-expressing oocytes in Kulori buffer at pH 6.0 containing either 20 μM carbonyl cyanide *m*-chlorophenyl hydrazone (H^+^ionophore, Sigma-Aldrich), 20 μM nigericin (K^+^/H^+^ exchanger, Abcam), or 20 μM valinomycin (K^+^ionophore, Abcam), for 15 min. Afterwards, we incubated the oocytes in Kulori buffer containing 100 μM I3M glucosinolate and the corresponding ionophore for 10 min. Assays performed with oocytes incubated in Kulori buffer without any ionophore served as a control.

The time course of I3M glucosinolate uptake was analyzed by incubating oocytes with 100 μM I3M glucosinolate in Kulori buffer (pH 6.0) for 5, 10, 20, 30, 45, 60, 120, 180, and 240 min, respectively. The apparent *Km* value of *Pa*GTR1 for I3M glucosinolate was determined by incubating injected oocytes in Kulori buffer (pH 6.0) for 10 min with different substrate concentrations. The *K*m value was calculated by nonlinear regression analysis in SigmaPlot 14.0 (Systat Software Inc.).

Each assay consisted of 14-15 oocytes and was stopped by washing oocytes four times with Kulori buffer. Afterwards, 12 of the washed oocytes were distributed into four Eppendorf tubes, with three oocytes per tube and immediately homogenized in 100 μL of 50% (v/v) methanol. After centrifugation (21,380 × *g* or 16, 000 × *g* for 15 min), the supernatant was incubated at −20 °C for at least 1 h to precipitate proteins, which were pelleted by centrifugation (21,380 × *g* or 16, 000 × *g* for 15 min). Finally, 60 μL sample was diluted with 120 μL ultrapure water, filtered through a 0.22 μm PVDF-based filter plate (Merck Millipore), and analyzed by LC-MS/MS. The glucosinolate concentration in oocytes was calculated by assuming an oocyte volume of 1 μL^63^.

### LC-MS/MS

Glucosinolates were quantified by LC-MS/MS using an Agilent 1200 HPLC system connected to an API3200 tandem mass spectrometer (AB SCIEX). Separation was achieved on an EC 250/4.6 NUCLEODUR Sphinx RP column (250 mm × 4.6 mm, 5 μm; Macherey–Nagel) using a binary solvent system consisting of 0.2% (v/v) formic acid in water (A) and acetonitrile (B), with a flow rate of 1 mL/min at 25 °C. The elution gradient was: 0-1 min, 1.5% B; 1-6 min, 1.5–5% B; 6-8 min, 5–7% B; 8-18 min, 7–21% B; 18-23 min, 21–29% B; 23-23.1 min, 29–100% B; 23.1-24 min, 100% B; 24-24.1 min, 100 to 1.5% B; 24.1-28 min, 1.5% B. Glucosinolates were detected in negative ionization mode. The ion spray voltage was set to −4,500 V. Gas temperature was set to 700 °C, curtain gas to 20 psi, collision gas to 10, nebulizing gas to 70 psi, and drying gas to 60 psi. Non-host glucosides were quantified using an Agilent 1200 HPLC system connected to an API5000 tandem mass spectrometer (AB SCIEX). Separation was achieved on an Agilent XDB-C18 column (5 cm × 4.6 mm, 1.8 μm) using a binary solvent system consisting of 0.05 % (v/v) formic acid in water (A) and acetonitrile (B) with a flow rate of 1.1 mL/min at 25 °C. The elution gradient was: 0-0.5 min, 5% B; 0.5-2.5 min, 5–31% B; 2.5-2.52 min, 31–100% B; 2.52-3.5 min, 100% B; 3.5-3.51 min, 100–5% B; 3.51-6 min, 5% B. Compounds were detected in negative ionization mode with ion spray voltage set to −4,500 V. The gas temperature was set to 700 °C, curtain gas to 30 psi, collision gas to 6, both nebulizing gas and drying gas to 60 psi. Multiple reaction monitoring (MRM) was used to monitor the transitions from precursor ion to product ion for each compound (Supplementary Table 7). Compounds were quantified using external standard curves. Analyst Software 1.6 Build 3773 (AB Sciex) was used for data acquisition and processing.

Samples from the pH-dependency experiment were analyzed by LC-MS/MS using an Advance UHPLC system (Bruker) connected to an EVOQ Elite TripleQuad mass spectrometer (Bruker) equipped with an electrospray ion source. Separation was achieved on an Kinetex 1.7u XB-C18 column (100 × 2.1 mm, 1.7 μm, 100 Å, Phenomenex) using a binary solvent system consisting of 0.05 % (v/v) formic acid in water (A) and acetonitrile with 0.05% (v/v) formic acid (B), with a flow rate of 0.4 mL/min at 40 °C. The elution gradient was: 0-0.2 min, 2% B; 0.2-1.8 min, 2-30% B; 1.8-2.5 min 30-100% B, 2.5-2.8 min 100% B, 2.8-2.9 min, 100-2% B and 2.9-4.0 min 2% B. Glucosinolates were detected in negative ionization mode. The instrument parameters were optimized by infusion experiments with pure standards. The ion spray voltage was set to −4,000 V. Cone temperature was set to 350 °C, cone gas to 20 psi, heated probe temperature to 200 °C, and probe gas flow to 50 psi. Nebulizing gas was set to 60 psi, and collision gas to 1.6 mTorr. MRM parameters are provided in Supplementary Table 7. Bruker MS Workstation software (Version 8.2.1, Bruker) was used for data acquisition and processing of glucosinolates. All other samples from experiments using oocyte expression system were analyzed using the LC-MS/MS method described above for insect cell-based assays. The concentrations of all glucosinolates in the substrate preference assay were determined using external standard curves. Assays performed to characterize I3M glucosinolate transport were quantified using 2-propenyl glucosinolate as internal standard.

### Double-stranded RNA synthesis

We synthesized seven different double-stranded RNAs (dsRNAs) between 120 and 298 bp in length, one specific for each *PaGTR1/2/3/5/6/7/8*, respectively, and a 223-bp fragment of the inducible metalloproteinase inhibitor (*IMPI*) from the greater wax moth *Galleria mellonella* (AY330624.1) (*dsIMPI*) using the T7 RiboMAX™ Express RNAi System (Promega). *In silico* off-target prediction was done by searches of all possible 21-mers of both RNA strands against the local *P. armoraciae* transcriptome database allowing for two mismatches. Except for *PaGTR5*, the prediction did not find any off-target towards putative transporter genes in the transcriptome (Supplementary Table 8).

### Function of *Pa*GTR1 *in vivo*

To analyze the function of *PaGTR1*, we injected newly emerged adult *P. armoraciae* beetles (reared on *B. juncea*) with 100 nL ultrapure water containing 80 ng of *dsPaGTR1* or 80 ng of *dsIMPI*, respectively, using a Nanoliter 2010 Injector (World Precision Instruments). Injected beetles were provided with detached leaves of three to four-week old *B. juncea* plants and kept in plastic containers with moistened tissue in the laboratory under ambient conditions. Four days after dsRNA injection, we collected *dsIMPI-injected* and *dsPaGTR1*-injected beetles for gene expression analysis (*n* = 5 replicates, three beetles per replicate) and glucosinolate analysis (*n* = 10 replicates, three beetles per replicate), respectively. The remaining beetles were used for a sequestration experiment with *Arabidopsis thaliana* Col-0 (*Arabidopsis*) plants that had been cultivated in a growth chamber at 21 °C, 55% relative humidity, and a 10-h photoperiod. To compare the accumulation and excretion of ingested glucosinolates in *dsIMPI*-injected and *dsPaGTR1*-injected beetles, we fed beetles with detached *Arabidopsis* leaves in Petri dishes (60 mm diameter) that contained 50 μL of ultrapure water and were sealed with parafilm to prevent leaf wilting (*n* = 9 replicates for day 4, *n* =10 replicates for other days, 5 beetles per replicate). Feeding assays were performed in the laboratory under ambient conditions, and leaves were exchanged every day for five consecutive days. To estimate how much the beetles fed, we determined the weight of each leaf before and after feeding. Feces were collected every day using 100 μL of ultrapure water per replicate, combined with 300 μL of pure methanol in a 1.5 mL Eppendorf tube and dried by vacuum centrifugation. Feces samples were then homogenized in 200 μL 80% (v/v) methanol using metal beads (2.4 mm diameter, Askubal) in a tissue lyzer (Qiagen) for 1 min at 30 Hz. After feeding, adults were starved for one day, weighed, frozen in liquid nitrogen, and stored at −20 °C until glucosinolate extraction. Beetle samples were homogenized using a plastic pestle in 200 μL 80% (v/v) methanol. All samples were then extracted with 1 mL 80% (v/v) methanol containing 25 μM 4-hydroxybenzyl glucosinolate as internal standard. After centrifugation (16,000 × *g* for 10 min), glucosinolates were extracted from the supernatant, converted to desulfo-glucosinolates, and analyzed by HPLC-UV as described before^23^. The glucosinolate content in adults or feces was calculated in nanomole per adult, respectively.

To confirm the specificity of *PaGTR1* knock-down we analyzed the effect of *dsPaGTR1* injection on the expression of *PaGTR2, PaGTR3, PaGTR9*, and *PaGTR10. PaGTR9* and *PaGTR10* share the highest nucleotide sequence similarity (69% sequence identity) with *PaGTR1*. *PaGTR2* and *PaGTR3* expression was analyzed because the recombinant transporters also used indol-3-ylmethyl (I3M) glucosinolate as a substrate. RNA extraction, purification, cDNA synthesis, and qPCR were performed as described before^61^.

### Function of *Pa*GTR5/6/7/8 *in vivo*

To analyze the function of *PaGTR5/6/7/8*, we injected newly emerged adult *P. armoraciae* beetles that had fed for two days on *B. juncea* leaves with 100 nL ultra pure water containing 100 ng of *dsIMPI* or each 100 ng of *dsPaGTR5/6/7/8* using a Nanoliter 2010 Injector (World Precision Instruments). A subset of the dsRNA-injected beetles were fed with detached leaves of *Arabidopsis* plants in plastic containers with moistened tissue for gene expression analysis (*n* = 6 replicates, two beetles per replicate). The remaining dsRNA-injected beetles were used for a sequestration experiment with *Arabidopsis* plants to compare the accumulation and excretion of previously stored and ingested glucosinolates in *dsIMPI*-injected and *dsPaGTR5/6/7/8*-injected beetles. We fed the injected beetles with detached *Arabidopsis* leaves in Petri dishes (60 mm diameter) that contained 30 μL of ultrapure water and were sealed with parafilm to prevent leaf wilting (*n* = 10 replicates, 6 beetles per replicate). Feeding assays were performed in the laboratory under ambient conditions, and leaves were exchanged every day for six consecutive days. To estimate how much the beetles fed, we determined the weight of each leaf before and after feeding. Starting from the second day, feces were collected as above for five days. After drying by vaccum centrifugation, feces were homogenized in 1 mL 80% (v/v) methanol using metal beads (2.4 mm diameter, Askubal) in a tissue lyzer (Qiagen) for 1 min at 30 Hz. Fed adults were starved for one day, weighed and frozen in liquid nitrogen until glucosinolate extraction. All samples were extracted and analyzed as described above except that an EC 250/4.6 NUCLEODUR 100-5 C18ec column (250 mm × 4.6 mm, 5 μm; Macherey–Nagel) was used for analyzing glucosinolates by HPLC-UV.

To confirm the specificity of *PaGTR5/6/7/8* knock-down, we analyzed the effect of *dsPaGTR5/6/7/8* injection on the expression of *PaGTR9* and *PaGTR10*, which share the highest nucleotide sequence similarity (67%-69% sequence identity) with *PaGTR5/6/7/8*. RNA extraction, purification, cDNA synthesis, and qPCR were performed as described before^61^.

### Function of *Pa*GTR2/3 *in vivo*

To analyze the functions of *PaGTR2* and *PaGTR3*, we injected third instar larvae of *P. armoraciae* (reared on *B. rapa*) with 100 nL ultrapure water containing 80 ng of *dsPaGTR2-*and 80 ng of *dsPaGTR3 (dsPaGTR2/3*) or 80 ng of *dsIMPI*, respectively, using a Nanoliter 2010 Injector (World Precision Instruments). Injected larvae were provided with detached *B. rapa* petioles and kept in plastic tubes with moistened tissue in the laboratory under ambient conditions until pupation. Newly emerged adults were injected with dsRNAs as described above and provided with detached *B. rapa* leaves. Three days after the second dsRNA injection, we collected *dsIMPI*-injected and *dsPaGTR2/3*-injected beetles for gene expression analysis (*n* = 6 replicates, two beetles per replicate) and glucosinolate analysis (*n* = 10 replicates, three beetles per replicate), respectively. The remaining beetles were used for a feeding experiment with *Arabidopsis*. Each replicate consisted of one *Arabidopsis* leaf and three beetles that were placed in a Petri dish (60 mm diameter) with 50 μL of ultrapure water (*n* = 13 replicates for *dsIMPI*-injected beetles, *n* = 12 replicates for *dsPaGTR2/3*-injected beetles). Each leaf was photographed before and after feeding to determine the consumed leaf area. Fed leaves were frozen in liquid nitrogen, freeze-dried, and homogenized using metal beads (2.4 mm diameter, Askubal) in a tissue lyzer (Qiagen) for 2 min at 30 Hz. Fed beetles were starved for one day, weighed, frozen in liquid nitrogen, and stored at −20 °C until glucosinolate extraction. Beetle samples were homogenized using a plastic pestle in 200 μL 80% (v/v) methanol. All samples were extracted with 1 mL 80% (v/v) methanol containing 25 μM 1-methylethyl glucosinolate (extracted from *Sisymbrium officinale* seeds) as internal standard. Glucosinolates were analysed after conversion to desulfo-glucosinolates by HPLC-UV as described before^23^. All the samples were additionally analyzed by another chromatographic run with the same method as above except that an EC 250/4.6 NUCLEODUR 100-5 C18ec column (250 mm × 4.6 mm, 5 μm; Macherey–Nagel) was used for analyzing glucosinolates that were not separated in the first chromatographic run. The glucosinolate content in adults and fed leaves was calculated in nanomole per adult and nanomole per cm^2^ leaf, respectively. The ingested glucosinolate amount was calculated based on the ingested leaf area and the corresponding leaf glucosinolate content. To elucidate which proportion of the ingested glucosinolates was sequestered, we expressed the glucosinolate amount detected in the beetles relative to the ingested glucosinolate amount from the leaves, which was set to 100%.

### Glucosinolate concentration in the hemolymph of adult *P. armoraciae*

Hemolymph was collected from seven-day old adult *P. armoraciae* reared on *B. juncea* by cutting off an abdominal leg and collecting the extruding droplet using glass capillaries (0.5 μL, Hirschmann^®^minicaps^®^) (*n* = 6 replicates, 50 beetles per replicate). The capillaries were marked with 1 mm intervals (corresponding to 15.6 nL) to estimate the volume of collected hemolymph. The hemolymph was diluted in 500 μL 90% (v/v) methanol, homogenized using metal beads (2.4 mm diameter, Askubal) in a tissue lyzer (Qiagen) for 1 min at 30 Hz, and boiled for 5 min at 95 °C. After two centrifugation steps (13,000 × *g* for 10 min each), the supernatant was dried by vacuum centrifugation, dissolved in 50 μL 50% (v/v) methanol, diluted in water, and analyzed by LC-MS/MS.

### Morphology of the Malpighian tubule system of *P. armoraciae*

To investigate the structure of the Malpighian tubule system, we dissected the gut including attached Malpighian tubules of four-day old *P. armoraciae* adults in PBS (pH 6.0) under a stereomicroscope. The tracheae that attach Malpighian tubules to the gut were removed using fine forceps to release the tubules. Pictures were taken with a Canon EOS 600D camera.

### pH of hemolymph and excretion fluid of isolated Malpighian tubules of *P. armoraciae*

The pH of the hemolymph and Malpighian tubule excretion fluid of five-day old *P. armoraciae* adults was assessed using the pH indicator bromothymol blue (Alfa Aesar). Hemolymph was collected by cutting off an abdominal leg and collecting the extruding droplet using a pipette (*n* = 3 replicates, six to ten beetles per replicate). Excretion fluid was collected from dissected Malpighian tubules that were incubated in saline A as described for the Ramsay assay (*n* = 4 replicates). Hemolymph and excretion fluid were immediately mixed with the same volume of 0.16% (w/v) bromothymol blue dissolved in 10% (v/v) ethanol, respectively, under water-saturated paraffin oil in a Sylgard-coated petri dish. The resulting color of the droplet was compared with those of citric acid–Na_2_HPO_4_ buffer solutions ranging from pH 5.2 to 6.6 in 0.2 increments mixed with 0.16% (w/v) bromothymol blue.

### Fate of plant glucosides injected in *P. armoraciae* beetles

To analyze the fate of plant glucosides *in vivo*,we injected 100 nL of an equimolar mixture of 2-propenyl glucosinolate, 4-hydroxybenzyl glucosinolate, linamarin, salicin and catalpol, each at 10 mM, and 0.15% (w/v) amaranth into the hemolymph of two-day old adult *P. armoraciae* (reared on *B. rapa*). One group of beetles was sampled 30 minutes after injection by freezing beetles in liquid nitrogen (*n* = 10 replicates, five beetles per replicate). The remaining beetles were fed with detached leaves of *Arabidopsis* in Petri dishes (60 mm diameter) in the laboratory under ambient conditions (*n* = 10 replicates, 5 beetles per replicate). We added 30 μL of ultrapure water to each Petri dish and sealed them with parafilm to prevent leaf wilting. After one day, we sampled the beetles as described above (*n* = 10 replicates, five beetles per replicate). Feces were collected using 100 μL of ultrapure water per replicate and combined with 300 μL of pure methanol in a 1.5 mL Eppendorf tube. All samples were stored at −20 °C until extraction. Beetle and feces samples were homogenized as described in the RNAi experiment. After centrifugation (16,000 × *g* for 10 min), the supernatant was dried by vacuum centrifugation, dissolved in 100 μL of ultrapure water and analyzed by LC-MS/MS. The glucoside content in adults or feces was calculated as nanomole per adult, respectively.

### Ramsay assay

To analyze the excretion of plant glucosides *in situ*, we performed Ramsay assays^64^ with dissected Malpighian tubules from four to five-day old *P. armoraciae* adults reared on *B. rapa* (Supplementary Fig. 6). Malpighian tubules were dissected in saline A (100 mM NaCl, 8.6 mM KCl, 2 mM CaCl_2_, 8.5 mM MgCl_2_, 4 mM NaH_2_PO_4_, 4 mM NaHCO_3_, 24 mM glucose, 10 mM proline, 25 mM 3-(N-Morpholino)propanesulfonic acid (MOPS), pH 6.0)^65^. Single tubules were transferred into a 10 μL droplet of saline B (60 mM NaCl, 10.3 mM KCl, 2.4 mM CaCl_2_, 10.2 mM MgCl_2_, 4.8 mM NaH_2_PO_4_, 4.8 mM NaHCO_3_, 28.8 mM glucose, 12 mM proline, 30 mM MOPS, 1 mM cyclic AMP, pH 6.0) under water-saturated paraffin oil in a Sylgard-coated petri dish. The proximal end of the tubule was pulled out of the droplet, attached to a metal pin and cut using a glass capillary to allow the collection of excretion fluid. To start the assay, we added two microliters of an equimolar glucoside mixture consisting of 2-propenyl glucosinolate, 4-methylsulfinylbutyl glucosinolate, 4-hydroxybenzyl glucosinolate, 2-phenylethyl glucosinolate, I3M glucosinolate, linamarin, salicin, catalpol, each at 40 mM, and 0.6% (w/v) amaranth) to saline B. After two to three hours, we collected the excretion fluid and 2 μL of the bathing droplet in 300 μL 80% (v/v) methanol, respectively. The Malpighian tubule was washed three times in about 15 mL of saline A and afterwards transferred into 300 μL of 80% (v/v) methanol. All samples were stored at −20 °C until extraction. Malpighian tubule samples were homogenized using a plastic pestle. After centrifugation (16,000 × *g* for 10 min), the supernatant was dried by vacuum centrifugation, dissolved in 70 μL ultrapure water and analyzed by LC-MS/MS. Out of 36 assays, 25 assays were excluded because no excretion fluid was visible within 2-3 hours.

### Statistical analysis

No statistical methods were used to predetermine sample size. Statistical analyses were conducted in R 3.5.1^66^ or in SigmaPlot 14.0 (Systat Software, Inc). Two groups were compared by two-tailed Student’s *t*-test, two-tailed Student’s paired *t*-test, Mann-Whitney *U* test or the method of generalized least squares^67^, depending on the variance homogeneity and the normality of residuals. Three or more groups were compared by one-way analysis of variance (ANOVA) followed by post hoc multiple comparisons test, or the method of generalized least squares^67^. If necessary, data were transformed prior to analysis. For comparisons using the method of generalized least squares, we applied the varIdent variance structure which allows each group to have a different variance. The *P*-value was obtained by comparing models with and without explanatory variable using a likelihood ratio test^68^. Significant differences between groups were revealed with factor level reductions^69^. Information about data transformation, statistical methods, and results of the statistical analyses are summarized in Supplementary Tables 3, 4 and 9.

## Reporting summary

Further information on research design is available in the Nature Research Reporting Summary linked to this article.

## Data availability

The data that support the findings of this study are available from the corresponding author upon request. Source data of this study is available at the open access data repository of the Max Planck Society (Edmond) under https://dx.doi.org/10.17617/3.4z. The sequences of the genes cloned in this paper have been deposited in the GenBank database (accession nos. MN433061–MN433082).

## Acknowledgements

We thank T. G. Köllner, D. G. Heckel, J. Gershenzon, S. O’Connor and M. Burow for comments on the manuscript, S. Donnerhacke, S. Gebauer-Jung, D. Schnabelrauch, L. Svenningsen, and A. Schilling for technical help, D. Yang for discussion on the project, and the greenhouse team at the MPI for Chemical Ecology for plant cultivation. This study was funded by the Max Planck Society. Additional support for this work was provided by Danish National Research Foundation grant DNRF99 (H. H. Nour-Eldin and C. Crocoll).

## Author contributions

Z.-L.Y., and F.B. designed the study. F.B. supervised the study. H.H.N. supervised the experiments with oocytes. Z.-L.Y. performed most of the experiments. S.H. contributed to RACE-PCR and gene cloning. M.R. and C.C. contributed to LC-MS/MS and HPLC-UV analyses. F.B. and H.V. performed transcriptome analysis. F.S. performed analysis of glucosinolate concentration in hemolymph. Z.-L.Y. and F.B. conducted data analysis. Z.-L.Y., and F.B. wrote the manuscript. All authors commented on the manuscript.

## Additional information

Correspondence and requests for materials should be addressed to F.B.

