## Supplementary Information for "Sugar transporters enable a leaf beetle to accumulate plant defense compounds"

**This file includes:**

Supplementary Figures 1-10

Supplementary Tables 1-9

**Additional Supplementary files for this manuscript include the following:**

Supplementary Data 1 (Excel Table)

Supplementary Data 2 (PDF Figure)

### Supplementary Figures 1-10

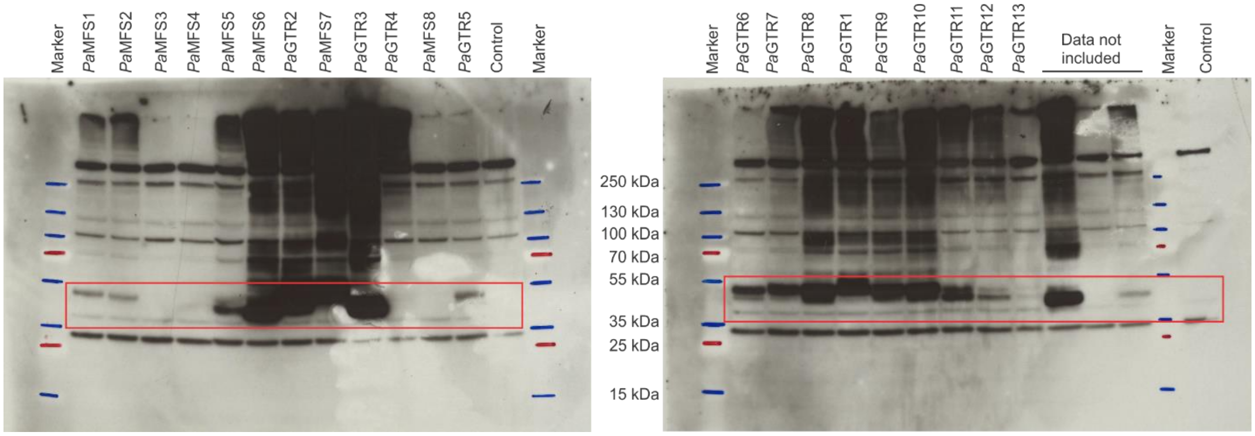

**Supplementary Figure 1. Detection of recombinant transporters expressed in High Five insect cells by Western blotting.** The region containing bands that correspond to recombinant proteins are framed in red. Protein marker bands were highlighted with blue and red lines on the film.

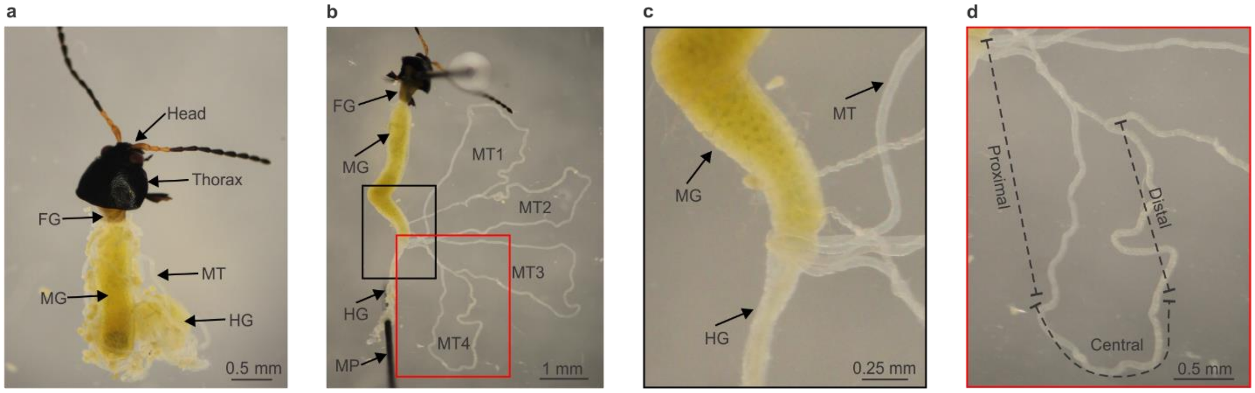

**Supplementary Figure 2. Morphology of Malpighian tubules. a** Gut with Malpighian tubules dissected from a four-day old *P. armoraciae* adult. **b** Malpighian tubule system consisting of four similar tubules (MT1-4) that empty at their proximal ends near the midgut-hindgut junction. Two tubules each fuse at their distal ends and appear to be attached to the midgut. **c** Magnification of the midgut-hindgut junction framed in black in panel **b**. **d** Magnification of one tubule (MT4) framed in red in panel **b**. FG, foregut; MG, midgut; HG, hindgut; MT, Malpighian tubules; MP, metal pin.

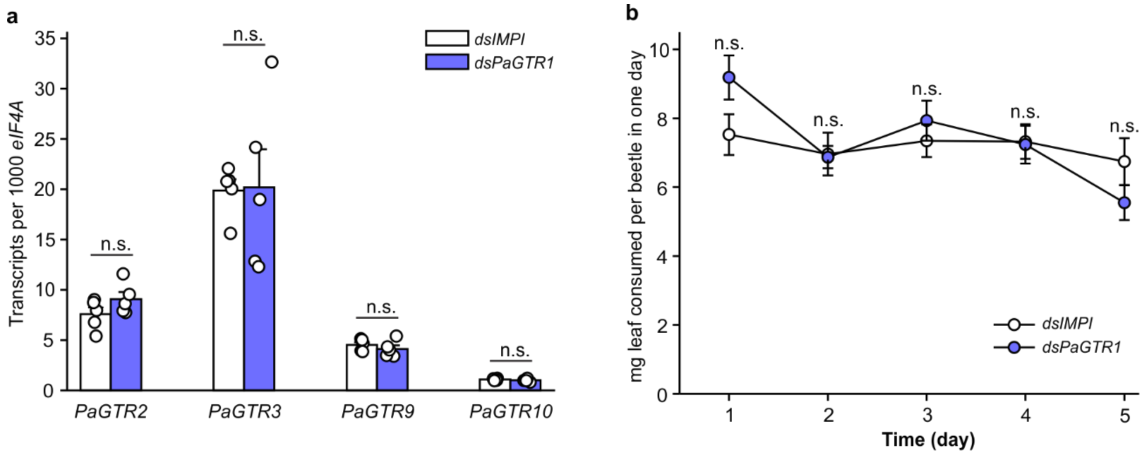

**Supplementary Figure 3. Analysis of potential off-target effects of *dsPaGTR1*-injection and comparison of beetle feeding on *Arabidopsis* leaves. a** Four days after dsRNA injection, the gene expression of *PaGTR2*, *PaGTR3*, *PaGTR9* and *PaGTR10* was determined by quantitative PCR to assess whether there was off-target silencing on other *PaGTR*s. The nucleotide sequence of *PaGTR1* was most similar to *PaGTR9* and *PaGTR10* in the *P. armoraciae* transcriptome. Upon heterologously expressed in High Five insect cells, *Pa*GTR2, *Pa*GTR3 and *Pa*GTR9 showed activity towards indol-3-ylmethyl glucosinolate ($n$ = 5). **b** Four days after dsRNA injection, adults were allowed to feed on *Arabidopsis* leaves for five days. Treatments were compared by two-tailed Student's $t$-test or Mann-Whitney $U$ test ($n$ = 10). Data are shown as mean ± s.e.m. n.s., not significantly different.

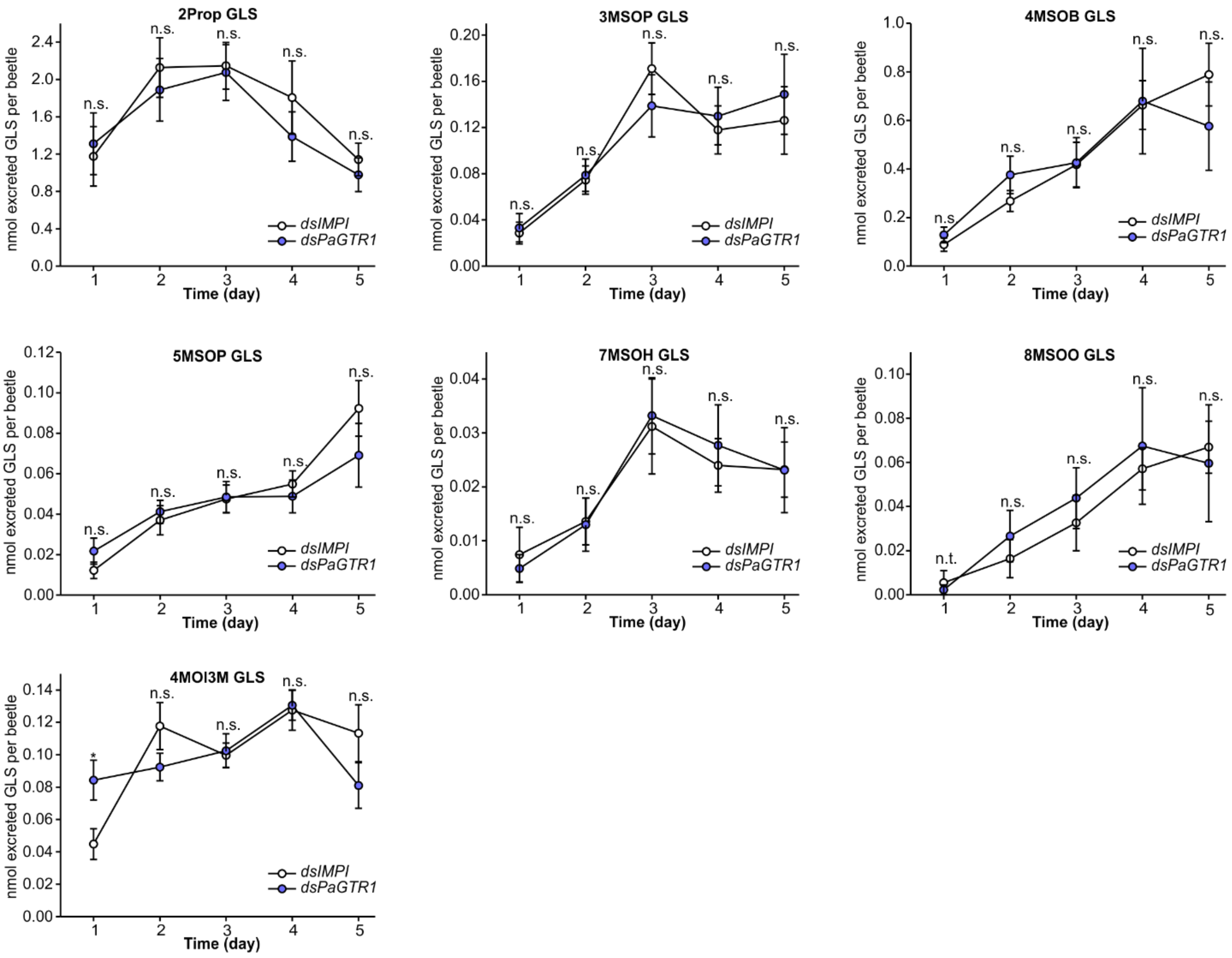

**Supplementary Figure 4. Time course of glucosinolate excretion during feeding on *Arabidopsis* by adult *P. armoraciae* after *dsIMPI*- or *dsPaGTR1*-injection.** The excreted glucosinolate amounts on each day were compared by two-tailed Student's *t*-test or Mann-Whitney *U* test ($n$ = 9 for day 4, $n$ =10 for other days). Data are shown as mean ± s.e.m. n.t., not tested; n.s., not significantly different; * $P < 0.05$; 2Prop, 2-propenyl; 3MSOP, 3-methylsulfinylpropyl; 4MSOB, 4-methylsulfinylbutyl; 5MSOP, 5-methylsulfinylpentyl; 7MSOH, 7-methylsulfinylheptyl; 4MOI3M, 4-methoxyindol-3-ylmethyl.

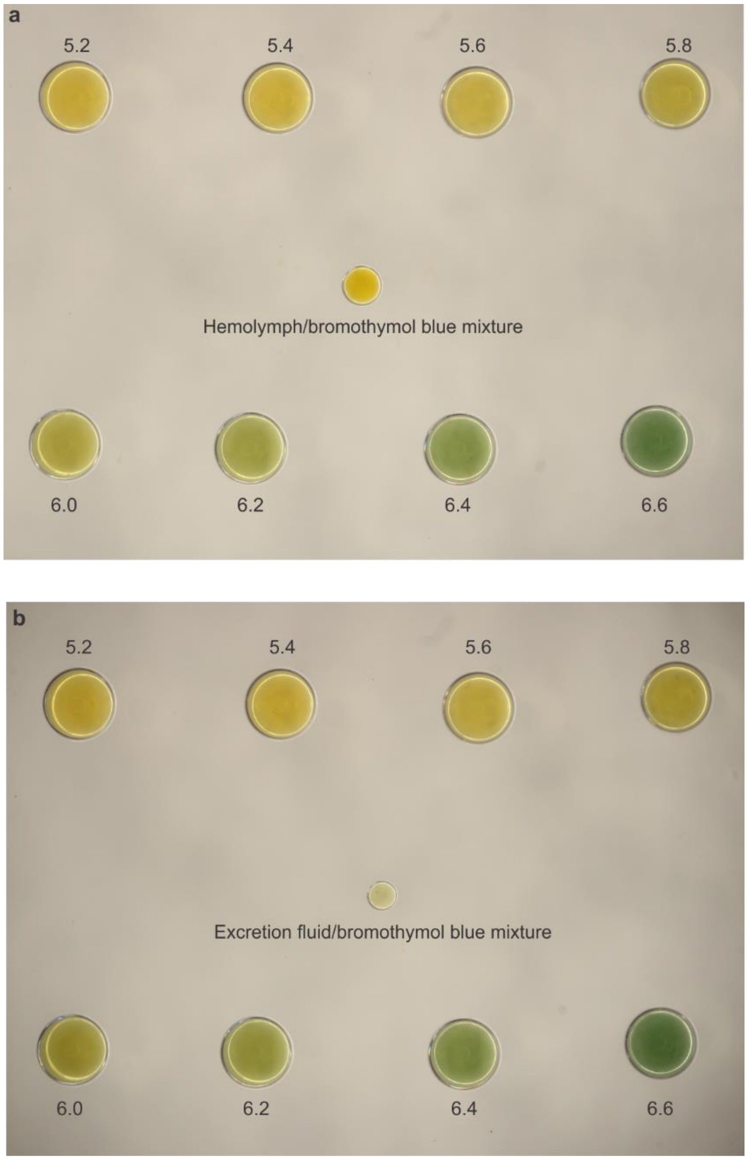

**Supplementary Figure 5. pH of hemolymph (a) and excretion fluid of isolated Malpighian tubules (b) of *P. armoraciae* adults.** Hemolymph and excretion fluid were mixed with an equal volume of 0.16% (w/v) bromothymol blue. Buffered standard solutions from pH 5.2 to 6.6 containing 0.08% bromothymol blue are shown as reference.

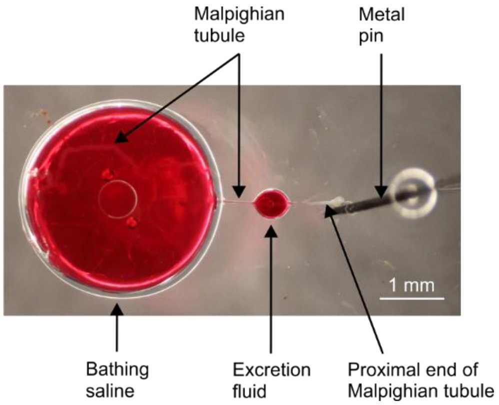

**Supplementary Figure 6. Preparation of *P. armoraciae* Malpighian tubule for glucoside excretion assay (Ramsay assay).** The dissected Malpighian tubule was placed in a droplet of bathing saline under water-saturated paraffin-oil, the proximal end was drawn out of the droplet and attached to the Sylgard-coated petri dish with a metal pin, and cut to allow the collection of excretion fluid. A mixture of eight different plant glucosides, each at a concentration of 6.7 mM, and 0.1% (w/v) amaranth was added to the saline. After 2-3 h, the bathing saline, Malpighian tubule and excretion fluid were sampled.

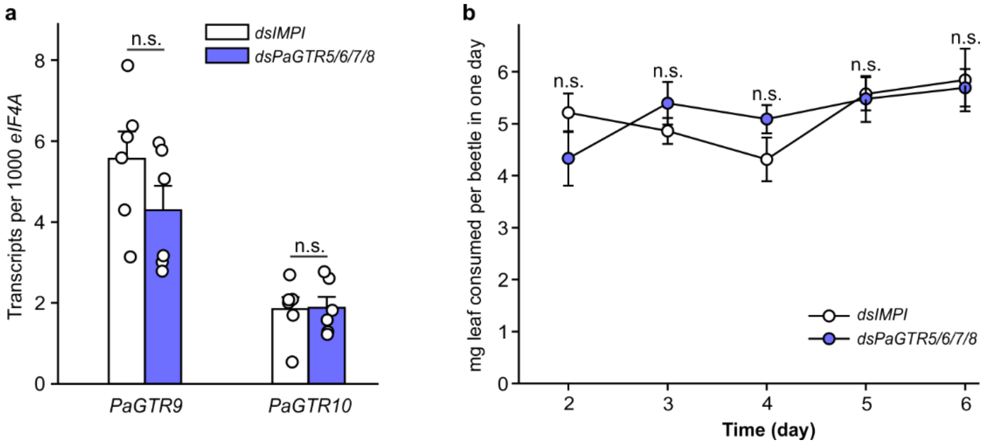

**Supplementary Figure 7. Analysis of potential off-target effects of *dsPaGTR5/6/7/8*-injection and comparison of beetle feeding on *Arabidopsis* leaves. a** Six days after dsRNA injection, the gene expression of *PaGTR9* and *PaGTR10* was determined by quantitative PCR to assess whether there was off-target silencing on other *PaGTR*s ($n$ = 6). The nucleotide sequence of *PaGTR5/6/7/8* was most similar to *PaGTR9* and *PaGTR10* in the *P. armoraciae* transcriptome. **b** After dsRNA injection, adults were allowed to feed on *Arabidopsis* leaves for six days. Feeding amount of each day was recorded from the second to the sixth day ($n$ = 10). Data are shown as mean ± s.e.m. n.s., not significantly different.

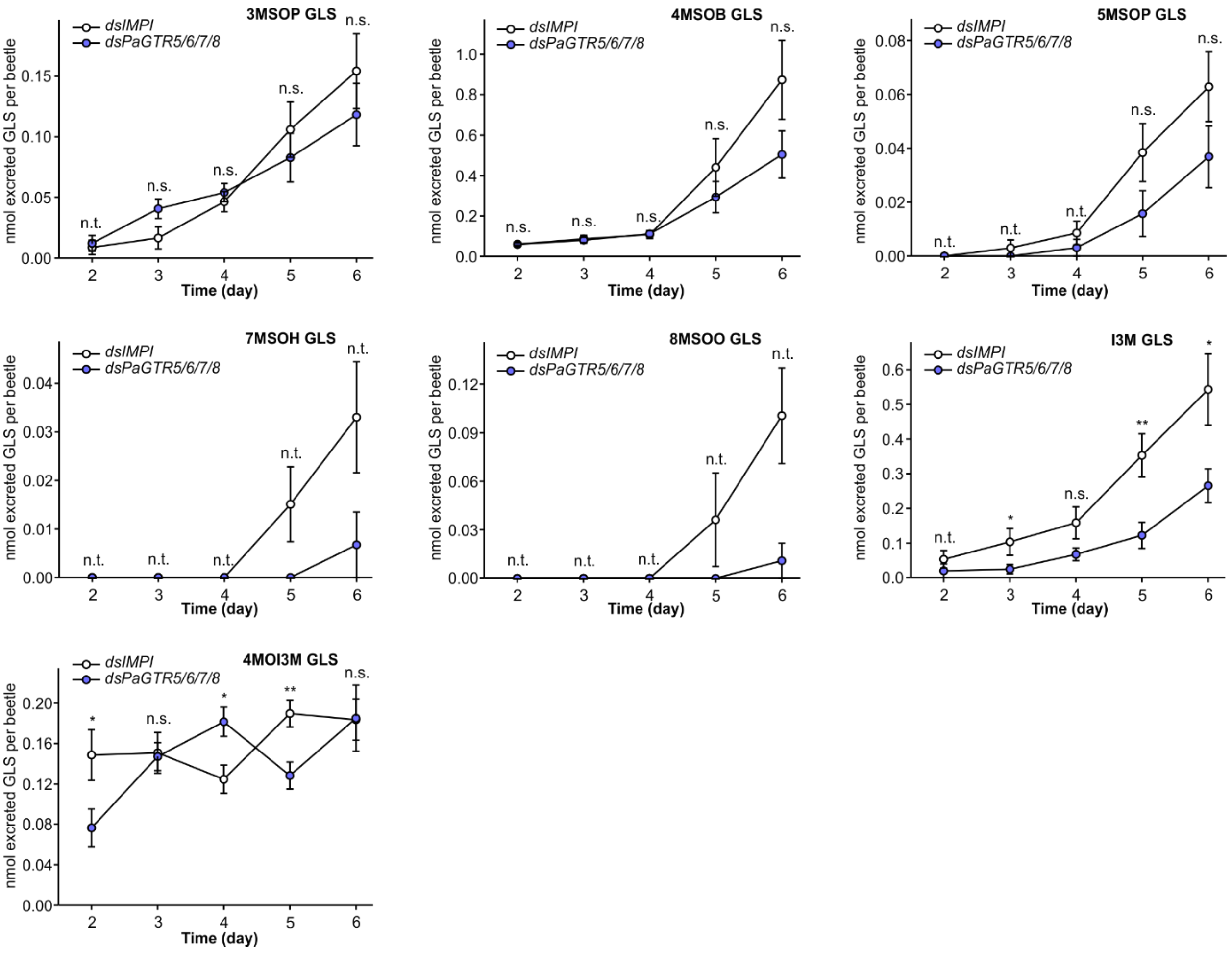

**Supplementary Figure 8. Time course of glucosinolate excretion during feeding on *Arabidopsis* by adult *P. armoraciae* after *dsIMPI*- or *dsPaGTR5/6/7/8*-injection.** The excreted glucosinolate amounts on each day were compared by two-tailed Student's *t*-test or Mann-Whitney *U* test ($n$ = 10). Data are shown as mean ± s.e.m. n.t., not tested; n.s., not significantly different; *$P$ < 0.05; **$P$ < 0.01; I3M, indol-3-ylmethyl.

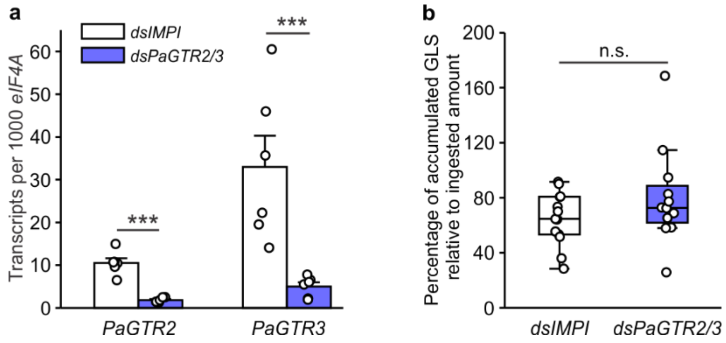

**Supplementary Figure 9. Accumulation of ingested glucosinolates in adult *P. armoraciae* beetles after knock-down of *PaGTR2* and *PaGTR3* expression. a** *PaGTR2 and PaGTR3* expression determined by quantitative PCR in adult *P. armoraciae* after injection of dsRNA targeting *PaGTR2* and *PaGTR3* (*dsPaGTR2/3*) or *IMPI* as a control ($n$ = 6). Data are shown as mean ± s.e.m. **b** Accumulation of the ingested glucosinolates in adult *P. armoraciae*. Three days after dsRNA-injection, adults were fed for one day with *Arabidopsis* leaves, starved for one day, and collected for glucosinolate extraction. Accumulated glucosinolates: sum of 3-methylsulfinylpropyl GLS, 3-methylthiopropyl GLS, 4-methylsulfinylbutyl GLS, 4-methylthiobutyl GLS, 7-methylsulfinylheptyl GLS and 8-methylsulfinyloctyl GLS. Due to a high background of indolic GLS, it was not possible to quantify the accumulation of ingested indolic GLS from *Arabidopsis*. Box plots show the median, interquartile range, and outliers of each data set ($n$ = 13 for the *dsIMPI*-injected beetles, $n$ = 12 for the *dsPaGTR2/3*-injected beetles). Treatments were compared by two-tailed Student's $t$-test. n.s., not significantly different; ***$P < 0.001$.

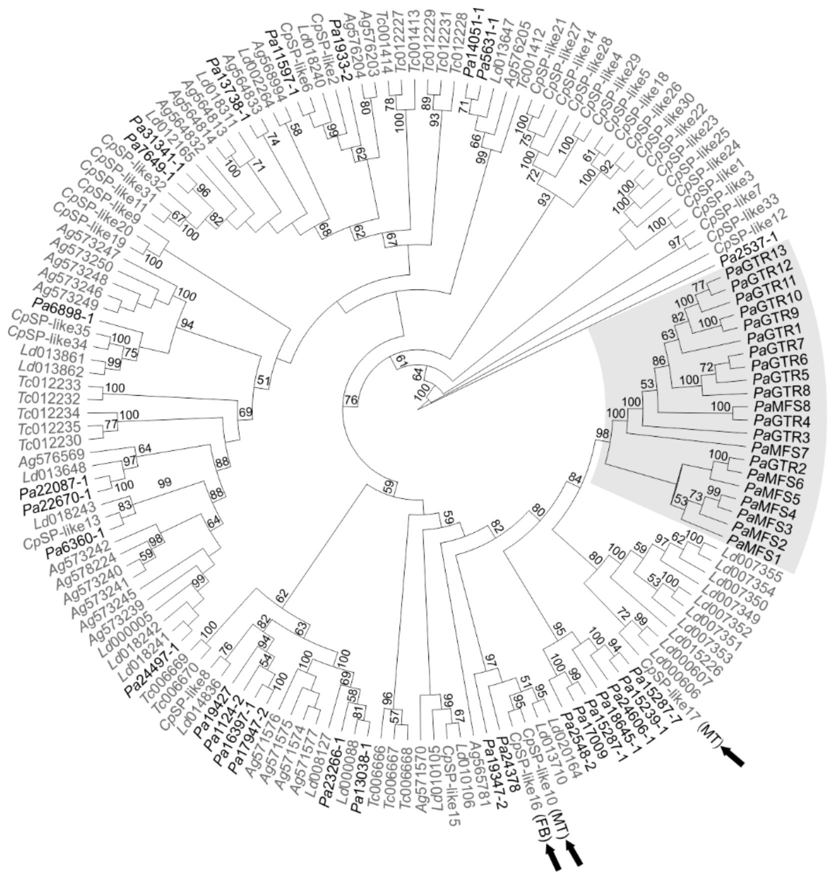

**Supplementary Figure 10. Diversification of coleopteran sugar porters.** Maximum-likelihood inferred phylogeny of a subset of predicted coleopteran sugar porters (Transporter Classification Database ID 2.A.1.1) identified in the *P. armoraciae* transcriptome (written in black) and the genomes of *Leptinotarsa decemlineata*, *Anoplophora glabripennis*, and *Tribolium castaneum* (marked with a black frame in Supplementary Data 2), and 35 sugar porters identified in the proteome of *Chrysomela populi* (written in grey). The *P. armoraciae*-specific clade investigated in this study is highlighted with a grey background. Tissue-specific localization of three sugar porters from *C. populi* that are closely related to *Pa*GTRs is written in parentheses. MT, Malpighian tubules; FB, fat body. Bootstrap support values higher than 50% are indicated on the corresponding branches. The tree was rooted with a putative vesicular neurotransmitter transporter (Transporter Classification Database ID 2.A.1.14) from *P. armoraciae* (*Pa*2537-1).

### Supplementary Tables 1-9

**Supplementary Table 1.** Numbers of putative transporters in *Phyllotreta armoraciae* transcriptome predicted by the Transporter Automatic Annotation Pipeline (TransAAP).

| **Transporter family** | **Transporter number** |
|---|---|
| The Major Facilitator Superfamily (MFS) | 353 |
| The ATP-binding Cassette (ABC) Superfamily | 174 |
| The H+- or Na+-translocating F-type, V-type and A-type ATPase (F-ATPase) Superfamily | 93 |
| The Voltage-gated Ion Channel (VIC) Superfamily | 71 |
| The Mitochondrial Carrier (MC) Family | 69 |
| The Neurotransmitter Receptor, Cys loop, Ligand-gated Ion Channel (LIC) Family | 39 |
| The P-type ATPase (P-ATPase) Superfamily | 38 |
| The HlyC/CorC (HCC) Family | 33 |
| The Epithelial Na+ Channel (ENaC) Family | 29 |
| The Drug/Metabolite Transporter (DMT) Superfamily | 26 |
| The Major Intrinsic Protein (MIP) Family | 26 |
| The Transient Receptor Potential Ca2+ Channel (TRP-CC) Family | 23 |
| The Amino Acid/Auxin Permease (AAAP) Family | 22 |
| The Glutamate-gated Ion Channel (GIC) Family of Neurotransmitter Receptors | 22 |
| The Neurotransmitter:Sodium Symporter (NSS) Family | 21 |
| The Amino Acid-Polyamine-Organocation (APC) Family | 18 |
| The Equilibrative Nucleoside Transporter (ENT) Family | 18 |
| The Mitochondrial Protein Translocase (MPT) Family | 17 |
| The Ca2+:Cation Antiporter (CaCA) Family | 14 |
| The Sulfate Permease (SulP) Family | 14 |
| The Ferrous Iron Uptake (FeoB) Family | 12 |
| The Resistance-Nodulation-Cell Division (RND) Superfamily | 12 |
| The YhaG Putative Tryptophan Uptake Permease (YhaG) family | 12 |
| The Unknown BART Superfamily-1 (UBS1) Family | 11 |

| | |
|---|---|
| The Solute:Sodium Symporter (SSS) Family | 10 |
| The Zinc (Zn2+)-Iron (Fe2+) Permease (ZIP) Family | 10 |
| The Choline Transporter Like (CTL) Family | 9 |
| The MerTP Mercuric Ion (Hg2+) Permease (MerTP) Family | 9 |
| The Multidrug/Oligosaccharidyl-lipid/Polysaccharide (MOP) Flippase Superfamily | 9 |
| The Chloride Carrier/Channel (ClC) Family | 8 |
| The Glycoside-Pentoside-Hexuronide (GPH):Cation Symporter Family | 8 |
| The Gap Junction-forming Innexin (Innexin) Family | 8 |
| The Annexin (Annexin) Family | 7 |
| The Cation-Chloride Cotransporter (CCC) Family | 7 |
| The Cation Diffusion Facilitator (CDF) Family | 7 |
| The Divalent Anion:Na+ Symporter (DASS) Family | 7 |
| The Organo Anion Transporter (OAT) Family | 7 |
| The Polycystin Cation Channel (PCC) Family | 6 |
| The Proton-dependent Oligopeptide Transporter (POT) Family | 6 |
| The Twin Arginine Targeting (Tat) Family | 6 |
| The Arsenite-Antimonite (ArsAB) Efflux Family | 5 |
| The Inward Rectifier K+ Channel (IRK-C) Family | 5 |
| The Ammonia Transporter Channel (Amt) Family | 4 |
| The Intracellular Chloride Channel (CLIC) Family | 4 |
| The Dicarboxylate/Amino Acid:Cation (Na+ or H+) Symporter (DAACS) Family | 4 |
| The Mg2+ Transporter-E (MgtE) Family | 4 |
| The Cytochrome Oxidase Biogenesis (Oxa1) Family | 4 |
| The Inorganic Phosphate Transporter (PiT) Family | 4 |
| The Anion Exchanger (AE) Family | 3 |
| The Arsenite-Antimonite (ArsB) Efflux Family | 3 |
| The ATP Exporter (ATP-E) Family | 3 |
| The Monovalent Cation:Proton Antiporter-1 (CPA1) Family | 3 |
| The Ca2+ Release-activated Ca2+ (CRAC) Channel (CRAC-C) Family | 3 |
| The Double Stranded RNA Transporter (dsRNA-T) Family | 3 |

| | |
|---|---|
| The H+-translocating Pyrophosphatase (H+-PPase) Family | 3 |
| The Iron/Lead Transporter (ILT) Superfamily | 3 |
| The Mitochondrial Tricarboxylate Carrier (MTC) Family | 3 |
| The Metal Ion (Mn2+-iron) Transporter (Nramp) Family | 3 |
| The Presenilin ER Ca2+ Leak Channel (Presenilin) Family | 3 |
| The ATP:ADP Antiporter (AAA) Family | 2 |
| The Anion Channel-forming Bestrophin (Bestrophin) Family | 2 |
| The Monovalent Cation:Proton Antiporter-2 (CPA2) Family | 2 |
| The Copper Transporter (Ctr) Family | 2 |
| The Type II (General) Secretory Pathway (IISP) Family | 2 |
| The Lysosomal Cystine Transporter (LCT) Family | 2 |
| The Branched Chain Amino Acid:Cation Symporter (LIVCS) Family | 2 |
| The CorA Metal Ion Transporter (MIT) Family | 2 |
| The Oligopeptide Transporter (OPT) Family | 2 |
| The 2-Hydroxycarboxylate Transporter (2-HCT) Family | 1 |
| The Auxin Efflux Carrier (AEC) Family | 1 |
| The Bile Acid:Na+ Symporter (BASS) Family | 1 |
| The Betaine/Carnitine/Choline Transporter (BCCT) Family | 1 |
| The Chloroplast Envelope Protein Translocase (CEPT or Tic-Toc) Family | 1 |
| The Concentrative Nucleoside Transporter (CNT) Family | 1 |
| The C4-Dicarboxylate Uptake (Dcu) Family | 1 |
| The C4-dicarboxylate Uptake C (DcuC) Family | 1 |
| The Epithelial Chloride Channel (E-ClC) Family | 1 |
| The Glutamate:Na+ Symporter (ESS) Family | 1 |
| The PTS Glucitol (Gut) Family | 1 |
| The Nucleotide-sensitive Anion-selective Channel, ICln (ICln) Family | 1 |
| The Magnesium Transporter1 (MagT1) Family | 1 |
| The 4 TMS Multidrug Endosomal Transporter (MET) Family | 1 |
| The H+- or Na+-translocating Bacterial Flagellar Motor 1ExbBD Outer Membrane Transport Energizer (Mot/Exb) Family | 1 |
| The Mitochondrial and Plastid Porin (MPP) Family | 1 |

| | |
|---|---|
| The Malonate:Na+ Symporter (MSS) Family | 1 |
| The NIPA Mg2+ Uptake Permease (NIPA) Family | 1 |
| The Non-selective Cation Channel-2 (NSCC2) Family | 1 |
| The OmpA-OmpF Porin (OOP) Family | 1 |
| The Reduced Folate Carrier (RFC) Family | 1 |
| The Telurite-resistance/Dicarboxylate Transporter (TDT) Family | 1 |
| The Homotrimeric Cation Channel (TRIC) Family | 1 |
| The Anion Channel Tweety (Tweety) Family | 1 |
| The Urea Transporter (UT) Family | 1 |

**Supplementary Table 2.** Glucosinolate concentrations in the hemolymph of seven-day old adult *P. armoraciae*.

| **Glucosinolate** | **Glucosinolate concentration (mM)** | | | | | | |
|---|---|---|---|---|---|---|---|
| | **Replicate 1** | **Replicate 2** | **Replicate 3** | **Replicate 4** | **Replicate 5** | **Replicate 6** | **Mean ± SD** |
| 2Prop GLS | 55.769 | 59.682 | 97.330 | 67.331 | 67.277 | 80.198 | 71.265 ± 15.263 |
| 3But GLS | 1.081 | 4.937 | 3.670 | 3.791 | 3.725 | 2.373 | 3.263 ± 1.343 |
| 2PE GLS | 0.157 | 0.302 | 0.938 | 0.704 | 0.280 | 0.342 | 0.454 ± 0.300 |
| Benzyl GLS | 0.345 | 0.386 | 0.634 | 0.735 | 0.401 | 0.258 | 0.460 ± 0.184 |
| I3M GLS | 1.948 | 1.811 | 3.118 | 2.626 | 1.289 | 1.702 | 2.082 ± 0.668 |
| Total | 59.300 | 67.118 | 105.689 | 75.187 | 72.972 | 84.873 | 77.523 ± 16.210 |

**Supplementary Table 3.** Glucosinolate levels in *PaGTR1*-silenced and control beetles.

| **Glucosinolate** | **nmol sequestered glucosinolate per adult (mean ± SD; *n* = 10)** | | **Statistical method[1]** | **Statistics** | ***P* value** |
|---|---|---|---|---|---|
| | ***dsIMPI*** | ***dsPaGTR1*** | | | |
| 3MSOP GLS | 1.231 ± 0.195 | 1.394 ± 0.282 | Two-tailed Student's *t*-test | *t* = -1.507 | 0.149 |
| 3MTP GLS | 0.188 ± 0.051 | 0.232 ± 0.088 | Two-tailed Student's *t*-test | *t* = -1.343 | 0.196 |
| 4MSOB GLS | 3.487 ± 0.491 | 4.087 ± 1.096 | Two-tailed Student's *t*-test | *t* = -1.581 | 0.131 |
| 4MTB GLS | 10.256 ± 2.100 | 11.200 ± 3.071 | Two-tailed Student's *t*-test | *t* = -0.803 | 0.433 |
| 5MSOP GLS | 0.191 ± 0.046 | 0.222 ± 0.032 | Two-tailed Student's *t*-test | *t* = -1.753 | 0.097 |
| 7MSOH GLS | 0.119 ± 0.032 | 0.158 ± 0.030 | Two-tailed Student's *t*-test | *t* = -2.830 | 0.011 |
| 8MSOO GLS | 1.135 ± 0.286 | 1.306 ± 0.441 | Two-tailed Student's *t*-test | *t* = -1.028 | 0.317 |
| 2Prop GLS | 55.841 ± 7.800 | 52.509 ± 3.984 | Mann-Whitney *U* test | *U* = 38.000 | 0.385 |
| 3-butenyl GLS | 0.935 ± 0.161 | 1.007± 0.295 | Two-tailed Student's *t*-test | *t* = -0.683 | 0.503 |
| I3M GLS | 2.830 ± 0.467 | 0.889 ± 0.303 | Two-tailed Student's *t*-test | *t* = 10.975 | < 0.001 |
| 4OHI3M GLS | 0.002 ± 0.004 | 0.002 ± 0.002 | Mann-Whitney *U* test | *U* = 50.000 | 1 |
| 4MOI3M GLS | 0.159 ± 0.084 | 0.227 ± 0.095 | Two-tailed Student's *t*-test | *t* = -1.687 | 0.109 |
| 1MOI3M GLS | 0.026 ± 0.008 | 0.004 ± 0.002 | Mann-Whitney *U* test | *U* = 0.000 | < 0.001 |
| Total | 76.399 ± 8.738 | 73.248 ± 4.350 | Two-tailed Student's *t*-test | *t* = 1.021 | 0.321 |

[1]Analyses were performed using SigmaPlot 14.0.

**Supplementary Table 4.** Glucosinolate levels in *PaGTR5/6/7/8*-silenced and control beetles.

| Glucosinolate | nmol sequestered glucosinolate per adult (mean ± SD; *n* = 10) | | Transformation | Statistics[1] | *P* value |
|---|---|---|---|---|---|
| | ***dsIMPI*** | ***dsPaGTR5/6/7/8*** | | | |
| 3MSOP GLS | 1.859 ± 0.217 | 1.713 ± 0.251 | - | *t* = 1.388 | 0.182 |
| 3MTP GLS | 0.432 ± 0.098 | 0.318 ± 0.087 | - | *t* = 2.739 | 0.013 |
| 4MSOB GLS | 4.555 ± 0.802 | 4.582 ± 0.855 | - | *t* = -0.073 | 0.943 |
| 4MTB GLS | 14.793 ± 1.872 | 12.287 ± 2.012 | - | *t* = 2.882 | 0.010 |
| 5MSOP GLS | 0.295 ± 0.048 | 0.315 ± 0.077 | Square-root | *t* = -0.648 | 0.525 |
| 7MSOH GLS | 0.218 ± 0.026 | 0.220 ± 0.045 | - | *t* = -0.139 | 0.891 |
| 8MSOO GLS | 1.317 ± 0.195 | 1.281 ± 0.376 | - | *t* = 0.263 | 0.795 |
| 2Prop GLS | 54.667 ± 5.272 | 47.919 ± 5.412 | - | *t* = 2.825 | 0.011 |
| 3-butenyl GLS | 1.287 ± 0.360 | 1.456 ± 0.584 | $Log_{10}$ | *t* = -0.791 | 0.439 |
| I3M GLS | 4.227 ± 0.487 | 5.225 ± 0.907 | - | *t* = -3.069 | 0.007 |
| 4MOI3M GLS | 0.301 ± 0.140 | 0.440 ± 0.173 | - | *t* = -1.977 | 0.064 |
| 1MOI3M GLS | 0.070 ± 0.010 | 0.070 ± 0.017 | - | *t* = -0.010 | 0.992 |
| Total | 84.021 ± 5.319 | 75.828 ± 4.321 | - | *t* = 3.781 | 0.001 |

[1]Analyses were performed with two-tailed Student's *t*-test using SigmaPlot 14.0.

**Supplementary Table 5.** Primers used in this study.

| Gene | Primer name | Primer sequence 5' - 3' | Use |
|---|---|---|---|
| *PaGTR1* | PaMFS28-CL1 | GTAAGTTTAAGTGTTATCTTCAAATTATCAAGGT | Cloning in pCR4-TOPO vector for sequencing; fwd |
| | PaMFS28-CL2 | CGATGAATGTGACGGAAGCA | Cloning in pCR4-TOPO vector for sequencing; rev |
| | PaMFS28-SEQ1 | ACAGAGGGAAATTCGGTTGT | Internal sequencing |
| | PaMFS28-SEQ2 | GCAGTTTCTGGTTCCGTTGT | Internal sequencing |
| | PaMFS28-IEX4-1 | ACGCGTCGACATGAACAATTGGACAAAAGAACATTTT | Expression of gene without stop codon in pIEx-4 vector; fwd |
| | PaMFS28-IEX4-2 | ATAAGAATGCGGCCGCGCCTTTTAATATCGCCTGAATTTC | Expression of gene without stop codon in pIEx-4 vector; rev |
| | PaMFS28-QPF1 | AGTTTCTGGTTCCGTTGTGC | qPCR; fwd |
| | PaMFS28-QPR1 | GGTTGCCCAATGATTCGGAT | qPCR; rev |
| | PaMFS28-NB1u-F1 | GGCTTAAUATGAACAATTGGACAAAAGAACATTTT | Cloning in pNB1u vector for protein expression in *Xenopus* oocytes; fwd |
| | PaMFS28-NB1u-R1 | GGTTTAAUTTAAGCGTAATCTGGAACATCGTATGGGTAGCCTTTTAATATCGCCTGAATTTC | Cloning in pNB1u vector for protein expression in *Xenopus* oocytes; rev |
| | T7-PaMFS28-F2 | TAATACGACTCACTATAGGGAGAGGTTTCGTTATTGGTCCATATTTCAGT | Amplification of DNA templates for dsRNA synthesis; fwd |
| | T7-PaMFS28-R2 | TAATACGACTCACTATAGGGAGAGCATTCGTCTTCTTTGCCCTTTT | Amplification of DNA templates for dsRNA synthesis; rev (141-bp amplified fragment of *PaGTR1* using the forward and reverse primers) |
| *PaGTR2* | c6623-1-CL1 | GCAGAAAGTGTCCGACAAATG | Cloning in pCR4-TOPO vector for sequencing; fwd |
| | c6623-1-CL2 | CATAACAACAATGTACAACTGTCGAG | Cloning in pCR4-TOPO vector for sequencing; rev |
| | c6623-1-EX1 | TGTCGACATGGTCAAAAAACGATACGAAATC | Expression of gene without stop codon in pIEx-4 vector; fwd |
| | c6623-1-EX2 | TGCGGCCGCATACTCTTTCAAAATCTTTTGTATTTCAAT | Expression of gene without stop codon in pIEx-4 vector; rev |
| | c6623-1-S1 | ACTTTCACGCAACCCACAAT | Internal sequencing |
| | c6623-1-S2 | GATTGGTGTTTGCCACGTT | Internal sequencing |
| | c6623-1-S3 | CAGCTTCGGCTATCCTTTGT | Internal sequencing |
| | C6623-1-QPF2 | GCCTGCGATACAACAACTGA | qPCR; fwd |

| | | | |
|---|---|---|---|
| | C6623-1-QPR2 | TCAAAGGAAGAGACCCGAGA | qPCR; rev |
| | T7-c6623-1-F1 | TAATACGACTCACTATAGGGAGAGGAAGCG AGAGTCGAAATTTC | Amplification of DNA templates for dsRNA synthesis; fwd |
| | T7-c6623-1-R1 | TAATACGACTCACTATAGGGAGAGCTAGAA GCAGCAGGGACAT | Amplification of DNA templates for dsRNA synthesis; rev (298-bp amplified fragment of *PaGTR2* using the forward and reverse primers) |
| *PaGTR3* | PaMFS27-CL1 | ACGGAATCTATGAATGATGTGCT | Cloning in pCR4-TOPO vector for sequencing; fwd |
| | PaMFS27-CL2 | AACCACCTGCATTTCGGAAC | Cloning in pCR4-TOPO vector for sequencing; rev |
| | PaMFS27-SEQ1 | AGCATCCTCTACATGACCCTG | Internal sequencing |
| | PaMFS27-SEQ2 | CTCGGTGGTGACGTCCAT | Internal sequencing |
| | PaMFS27-SEQ3 | GGGATTTCTACTCATGGTGATCT | Internal sequencing |
| | PaMFS27-IEX4-1 | ACGCGTCGACATGAAGGACGAGATATCTCT GAAG | Expression of gene without stop codon in pIEx-4 vector; fwd |
| | PaMFS27-IEX4-2 | ATAAGAATGCGGCCGCATAACTCTTGAGCT CCGCCT | Expression of gene without stop codon in pIEx-4 vector; rev |
| | PaMFS27-QPF1 | CGCGTTGTTATTGGGATGCT | qPCR; fwd |
| | PaMFS27-QPR1 | GCTTGTGTTTGCTTGGTCGA | qPCR; rev |
| | T7-PaMFS27-F2 | TAATACGACTCACTATAGGGAGATATCCTC CACAGCCACAGGAAT | Amplification of DNA templates for dsRNA synthesis; fwd |
| | T7-PaMFS27-R2 | TAATACGACTCACTATAGGGAGAGAACAGC TCGGTGACCAGCA | Amplification of DNA templates for dsRNA synthesis; rev (182-bp amplified fragment of *PaGTR3* using the forward and reverse primers) |
| *PaGTR4* | PaMFS43-CL1 | TACCGTCTGCCTCATACTCG | Cloning in pCR4-TOPO vector for sequencing; fwd |
| | PaMFS43-CL2 | TGCCTGACTAGTCTACACCA | Cloning in pCR4-TOPO vector for sequencing; rev |
| | PaMFS43-SEQ1 | TGAGCGTTGTACCTGTATACAT | Internal sequencing |
| | PaMFS43-IEX4-1 | ACGCGTCGACATGAAGGTAAAACAAGATG AAGATTACC | Expression of gene without stop codon in pIEx-4 vector; fwd |
| | PaMFS43-IEX4-2 | ATAAGAATGCGGCCGCTCTTTTAAGTAATTT TTCTATTTCACTGTAATT | Expression of gene without stop codon in pIEx-4 vector; rev |
| | PaMFS43-QPF2 | GTTACACGGGAATGCTGCAT | qPCR; fwd |
| | PaMFS43-QPR2 | CGGGCATCAAAGGAAACACA | qPCR; rev |
| *PaGTR5* | PaMFS15_3R_a | TACGCAGCTGAAGTTAGCGAGGATCACA | 3' RACE |
| | PaMFS15_3R_b | AGCACCGTTTTTAGACCAAGCAGGCACT | 3' RACE |

| | | | |
|---|---|---|---|
| | PaMFS15-5R | TATAGCGGTACCAGTGGAAGATATAACAAGCA | 5 'RACE |
| | PaMFS15-CL1 | GTAAAAGGTGCGACAAAACTGT | Cloning in pCR4-TOPO vector for sequencing; fwd |
| | PaMFS15-CL2 | TACTGCACATTATTAAAATTGACAATT | Cloning in pCR4-TOPO vector for sequencing; rev |
| | PaMFS15-SEQ1 | TTCCCGGAGCTTGGATAGTT | Internal sequencing |
| | PaMFS15-IEX4-1 | ACGCGTCGACATGGATAACAAAAAATATGAGAATATTCAAAAG | Expression of gene without stop codon in pIEx-4 vector; fwd |
| | PaMFS15-IEX4-2 | ATAAGAATGCGGCCGCTTTTTTCGCATAATTTCTCAATATTTCCT | Expression of gene without stop codon in pIEx-4 vector; rev |
| | PaMFS15-QPF1 | GGTTTAGGTCCTATTCCACTAGCT | qPCR; fwd |
| | PaMFS15-QPR1 | GTTACCCAAAGATTCCGACACTAG | qPCR; rev |
| | T7-PaMFS15-F6 | TAATACGACTCACTATAGGGAGAACAGTTTGCTGCTTTGGATCC | Amplification of DNA templates for dsRNA synthesis; fwd |
| | T7-PaMFS15-R6 | TAATACGACTCACTATAGGGAGATACCCAAAGATTCCGACACTAGG | Amplification of DNA templates for dsRNA synthesis; rev (169-bp amplified fragment of *PaGTR5* using the forward and reverse primers) |
| *PaGTR6* | PaMFS41_5R_a | AGTCACCGCTCTGAGCTCTGGTAATGGT | 5' RACE |
| | PaMFS41_5R_b | ACGCCTGAAAGCAGCATTACAAGCCCTGA | 5 'RACE |
| | PaMFS41-CL1 | CGTCTAAATCAAGTGAAATTGTATCGAAT | Cloning in pCR4-TOPO vector for sequencing; fwd |
| | PaMFS41-CL2 | GTGAGGGTCAATTTGTAATATAAATCGT | Cloning in pCR4-TOPO vector for sequencing; rev |
| | PaMFS41-SEQ1 | CAATAGAGGAAAATTCGGTTGTTATTTC | Internal sequencing |
| | PaMFS41-IEX4-1 | ACGCGTCGACATGGATAACCAAAATGAGAATAATCAAAA | Expression of gene without stop codon in pIEx-4 vector; fwd |
| | PaMFS41-IEX4-2 | ATAAGAATGCGGCCGCAATATTCTTCGAGACATCATAGTTTTTCA | Expression of gene without stop codon in pIEx-4 vector; rev |
| | PaMFS41-QPF3 | TCCAACAAATTGGGCATCTAT | qPCR; fwd |
| | PaMFS41-QPR3 | AACGAATACCAACAAAGGAGAAGT | qPCR; rev |
| | T7-PaMFS41-F1 | TAATACGACTCACTATAGGGAGATGGATTAGGTTTAGGTCCAATTG | Amplification of DNA templates for dsRNA synthesis; fwd |
| | T7-PaMFS41-R1 | TAATACGACTCACTATAGGGAGAGTTAATGCAAATCCTACAACACTACC | Amplification of DNA templates for dsRNA synthesis; rev (120-bp amplified fragment of *PaGTR6* using the forward and reverse primers) |
| *PaGTR7* | PaMFS40_3R | TATTCAGGGCTCGTAATGCTGCTGGCTA | 3' RACE |
| | PaMFS40-CL1 | AGGTGCATTAGAATTTTTAATTGGATTG | Cloning in pCR4-TOPO vector for sequencing; fwd |

| | | | |
|---|---|---|---|
| | PaMFS40-CL2 | CCAAGAAGTCCATAACTATTAAGATAGACT | Cloning in pCR4-TOPO vector for sequencing; rev |
| | PaMFS40_SEQ1 | ACACCAATACATTTGTTGGTTAA | Internal sequencing |
| | PaMFS40_SEQ2 | TATTACAGTAACACCCGAGCAGT | Internal sequencing |
| | PaMFS40-IEX4-1 | ACGCGTCGACATGGATAACCAAACTAAGAA AAATCAAAA | Expression of gene without stop codon in pIEx-4 vector; fwd |
| | PaMFS40-IEX4-2 | ATAAGAATGCGGCCGCTTTTTTCGCATAATT TCTCAATATTTCCT | Expression of gene without stop codon in pIEx-4 vector; rev |
| | PaMFS40-QPF1 | CCAAAGGAAGCATAAGGAGCAT | qPCR; fwd |
| | PaMFS40-QPR1 | CAGTAACACCCGAGCAGTATTTAT | qPCR; rev |
| | T7-PaMFS40-F1 | TAATACGACTCACTATAGGGAGAT TACAGTTTGCTTCTTTGGATTGG | Amplification of DNA templates for dsRNA synthesis; fwd |
| | T7-PaMFS40-R1 | TAATACGACTCACTATAGGGAGAG ATATAAAAATGTTAACGTAGAAGATGCAA | Amplification of DNA templates for dsRNA synthesis; rev (147-bp amplified fragment of *PaGTR7* using the forward and reverse primers) |
| *PaGTR8* | PaMFS48_3R | ACCGCGCCTGCGTTGTTATTCGTTGTTA | 3' RACE |
| | PaMFS48_5R | AGTCTCCCGATAAGCTGGTGCCTGCT | 5' RACE |
| | PaMFS48-CL1 | GCTGAGCAAATAAAGTGTCGAATT | Cloning in pCR4-TOPO vector for sequencing; fwd |
| | PaMFS48-CL2 | ATGTATATTTTTGTAACTAATAAGAGCCAA AA | Cloning in pCR4-TOPO vector for sequencing; rev |
| | PaMFS48-QPF1 | CTAATGCCCAAACTCTGCGAC | Internal sequencing |
| | PaMFS48-IEX4-1 | ACGCGTCGACATGAGAAAAATGAGTTTAGA AGACCAT | Expression of gene without stop codon in pIEx-4 vector; fwd |
| | PaMFS48-IEX4-2 | ATAAGAATGCGGCCGCATTTTTCACGTTACT TTTCAACATTTC | Expression of gene without stop codon in pIEx-4 vector; rev |
| | PaMFS48-QPF4 | GGTTTAATACCGTTGCTGACG | qPCR; fwd |
| | PaMFS48-QPR4 | CCTACCGAATCTCTCGATAACG | qPCR; rev |
| | T7-PaMFS48-F1 | TAATACGACTCACTATAGGGAGAT CAATTCATCATTATTGGTACAACTC | Amplification of DNA templates for dsRNA synthesis; fwd |
| | T7-PaMFS48-R1 | TAATACGACTCACTATAGGGAGAA AGATGCAGAACTGCCGATT | Amplification of DNA templates for dsRNA synthesis; rev (180-bp amplified fragment of *PaGTR8* using the forward and reverse primers) |
| *PaGTR9* | PaMFS16_5R_a | TGCGCCACTGAAACCTAAACCTACGCCT | 5' RACE |
| | PaMFS16_5R_b | ACACCGTAACCGCCCTGATCTCGGCT | 5' RACE |
| | PaMFS16-CL3 | AATGGTGCTTGCAGTGGTTT | Cloning in pCR4-TOPO vector for sequencing; fwd |

| | | | |
|---|---|---|---|
| | PaMFS16-CL4 | CGCGTAAAATCGGCTTAATATGA | Cloning in pCR4-TOPO vector for sequencing; rev |
| | PaMFS16-IEX4-3 | GTCGACATGGATTTAGAAAATAAAACACATCAAACA | Expression of gene without stop codon in pIEx-4 vector; fwd |
| | PaMFS16-IEX4-4 | GCGGCCGCTTTTATTCGAAGCTTCGAATAATTTAACA | Expression of gene without stop codon in pIEx-4 vector; rev |
| | PaMFS16-QPF1 | CATCGTTAATAGCGTACCTTCAATG | qPCR; fwd |
| | PaMFS16-QPR1 | CAATTCAGAGACCCACGCTTG | qPCR; rev, internal sequencing |
| *PaGTR10* | PaMFS42_5R | ACCATGTCGCCTGATAAGCTAGTGCCTGCT | 5' RACE |
| | PaMFS42-CL5 | GACGATACAAAATTTGAAAACTCAATAAATAAG | Cloning in pCR4-TOPO vector for sequencing; fwd |
| | PaMFS42-CL8 | CATTCTAACATTATGGCATTTAAAAGTATAAGT | Cloning in pCR4-TOPO vector for sequencing; rev |
| | PaMFS42-SEQ1 | CTGGTTGGGTGGTTTTCTCG | Internal sequencing |
| | PaMFS42-IEX4-1 | ACGCGTCGACATGGATTTAGAAAATAAGCCGCAT | Expression of gene without stop codon in pIEx-4 vector; fwd |
| | PaMFS42-IEX4-3 | ATAAGAATGCGGCCGCTTTTATTTGTAGCTTCGAATAATTTTTTAGC | Expression of gene without stop codon in pIEx-4 vector; rev |
| | PaMFS42-QPF1 | TCATCGTTAATAGCGTACCTTCAATG | qPCR; fwd |
| | PaMFS42-QPR1 | GCAGTTCGGACGTCATCACTT | qPCR; rev |
| *PaGTR11* | PaMFS45_3R | CACTGGGAGTATCAATGGCTTGGACATC | 3' RACE |
| | PaMFS45_3Rb | AGAGATGGAAGAGAGTCACAACGTTGCGA | 3' RACE |
| | PaMFS45_3Rc | ACCGAGGCTTATTCCGTGATGTCGATATACGT | 3' RACE |
| | PaMFS45_5R | CGGACCGATAATGTAACCAAGAAGATGCCCCA | 5' RACE |
| | PaMFS45-CL3 | CATTAAGAAGCACCAACTACCGTAG | Cloning in pCR4-TOPO vector for sequencing; fwd |
| | PaMFS45-CL4 | CGGTTGGTTATATTGGTTAAATTTACAA | Cloning in pCR4-TOPO vector for sequencing; rev |
| | PaMFS45-SEQ1 | CAAATGGGGCATCTTCTTGGT | Internal sequencing |
| | PaMFS45-IEX4-1 | ACGCGTCGACATGGAAGAGAGTCACAACGTT | Expression of gene without stop codon in pIEx-4 vector; fwd |
| | PaMFS45-IEX4-2 | ATAAGAATGCGGCCGCATGTATTTTAAGTTCGGAATAATTTTTCAAC | Expression of gene without stop codon in pIEx-4 vector; rev |
| | PaMFS45-QPF1 | TCACTGGGAGTATCAATGGCTT | qPCR; fwd |
| | PaMFS45-QPR1 | ACACCGCAAAATAACAAATATCTCTTC | qPCR; rev |

| | | | |
|---|---|---|---|
| *PaGTR12* | PaMFS14_3R_a | TGACTCTTTCACTGGGAGCTTCAATGTCTTGGA | 3' RACE |
| | PaMFS14_3R_b | TTCTCAATGGTGCCGGAAACACCCATATATTTG | 3' RACE |
| | PaMFS14-CL1 | AAAGAAAACCAAAAAAATTAAAATTTTTTTAAGAAA | Cloning in pCR4-TOPO vector for sequencing; fwd |
| | PaMFS14-CL2 | CTAATTCATATTAACACAGCAAACTTAGT | Cloning in pCR4-TOPO vector for sequencing; rev |
| | PaMFS14_SEQ1 | ACCGAAGCTTATTCCGTAACGT | Internal sequencing |
| | PaMFS14-IEX4-3 | ACGCGTCGACATGCATTTAGAGATAAAAGAAAACCACAAAG | Expression of gene without stop codon in pIEx-4 vector; fwd |
| | PaMFS14-IEX4-2 | ATAAGAATGCGGCCGCACGTATTTTAAGTTCGGAATAATTTTTTAAC | Expression of gene without stop codon in pIEx-4 vector; rev |
| | PaMFS14-QPF1 | GTTTTCAAAGTAACTTCCTACGTTCT | qPCR; fwd |
| | PaMFS14-QPR1 | TATGAAGAACAAAGATAGGGCAGTAC | qPCR; rev |
| *PaGTR13* | PaMFS46_3R_a | GAGCATCCCTAACTTGGACATCTCCAGT | 3' RACE |
| | PaMFS46_5R_a | TATCGATGTCACGGAATGAGCCTCGGTA | 5' RACE |
| | PaMFS46_5R_b | AATATATAGGTGTTTCCGGCGCCATTGAGAAA | 5' RACE |
| | PaMFS46-CL1 | GGCATTAAAGAACATTTAAGAAGTACCAA | Cloning in pCR4-TOPO vector for sequencing; fwd |
| | PaMFS46-CL2 | CAACATTTAGAATGTCTGGAGATTTTTTAG | Cloning in pCR4-TOPO vector for sequencing; rev |
| | PaMFS46_SEQ1 | TCGATATTCCATCAAATGGGACA | Internal sequencing |
| | PaMFS46-IEX4-1 | ACGCGTCGACATGCATTTAGAGATGGAAGAAAGTC | Expression of gene without stop codon in pIEx-4 vector; fwd |
| | PaMFS46-IEX4-2 | ATAAGAATGCGGCCGCATGTATTTTAAGTTGGGAATAATTTTTTAACAT | Expression of gene without stop codon in pIEx-4 vector; rev |
| | PaMFS46-QPF3 | GTTTAGGACCTATTCCGCACAT | qPCR; fwd |
| | PaMFS46-QPR3 | GAAAACAAACACTACAACGTAACAGG | qPCR; rev |
| *PaMFS1* | Pa3391-CL1 | CATTCTCCTAATCTTGGCATTATCTAT | Cloning in pCR4-TOPO vector for sequencing; fwd |
| | Pa3391-CL2 | GTCTGCCTGCAGAAGATTTTAC | Cloning in pCR4-TOPO vector for sequencing; rev |
| | Pa3391-EX1 | TGTCGACATGAGTTCTGGCATAGATCAAAAAT | Expression of gene without stop codon in pIEx-4 vector; fwd |
| | Pa3391-EX2 | TGCGGCCGCCCTGGTTTTCAGGTATTCCTG | Expression of gene without stop codon in pIEx-4 vector; rev |
| | Pa3391-S1 | GTGGGTCGGAAGAGATGCTT | Internal sequencing |

| | | | |
|---|---|---|---|
| | Pa3391-QPF1 | CAATTCGGCAGGTACCAGTT | qPCR; fwd |
| | Pa3391-QPR1 | TTCGTCTCCTGCCAGTCTTT | qPCR; rev |
| *PaMFS2* | Pa6623-7-CL1 | GAGCAATTACTTAATTAGATGGCAGT | Cloning in pCR4-TOPO vector for sequencing; fwd |
| | Pa6623-7-CL2 | ATACACACAATGTACCACAGCTTAAA | Cloning in pCR4-TOPO vector for sequencing; rev |
| | Pa6623-7-EX1 | TGTCGACATGGTCGAAATGGGCAAAG | Expression of gene without stop codon in pIEx-4 vector; fwd |
| | Pa6623-7-EX2 | TGCGGCCGCATTTCCAATCTCCTCCTGGATT | Expression of gene without stop codon in pIEx-4 vector; rev |
| | Pa6623-7-S1 | AGCGTTATCATGGCTGGATT | Internal sequencing |
| | Pa6623-7-QPF2 | ACAGTACATGAGCGGTGCAG | qPCR; fwd |
| | Pa6623-7-QPR2 | GCGCTACACTATCGCCAGAT | qPCR; rev |
| *PaMFS3* | Pa19756-1-CL1 | GTTGAGGTGATTGCAGGAAC | Cloning in pCR4-TOPO vector for sequencing; fwd |
| | Pa19756-1-CL2 | GTTAGACTGATTGCACTTGAGGA | Cloning in pCR4-TOPO vector for sequencing; rev |
| | Pa19756-1-EX1 | TGTCGACATGGGTAAATTGGAAGAAGTAAA ATATG | Expression of gene without stop codon in pIEx-4 vector; fwd |
| | Pa19756-1-EX2 | TGCGGCCGCAGCGCATTTTCTAAACAATTG AA | Expression of gene without stop codon in pIEx-4 vector; rev |
| | Pa19756-1-S1 | CTGACGGAACTGTCGGAGAAT | Internal sequencing |
| | Pa19756-1-QPF2 | TGCAGTGTCTTTCTCCGTTG | qPCR; fwd |
| | Pa19756-1-QPR2 | CCGTAAACAATCCCACGAAC | qPCR; rev |
| *PaMFS4* | Pa20117-CL1 | CACCCTGCATTGTATCAGGATT | Cloning in pCR4-TOPO vector for sequencing; fwd |
| | Pa20117-CL2 | TTTACTTAAAACTTGACAAACATGACTT | Cloning in pCR4-TOPO vector for sequencing; rev |
| | Pa20117-EX1 | TGTCGACATGCTCTCCGGAAGCGCC | Expression of gene without stop codon in pIEx-4 vector; fwd |
| | Pa20117-EX2 | TGCGGCCGCCCAACGTTTCAATTTAATTCGA TA | Expression of gene without stop codon in pIEx-4 vector; rev |
| | Pa20117-S1 | GAGCTACAAGCCTTTCACGTT | Internal sequencing |
| | Pa20117-QPF4 | CACTGCGATAGTCGGTGTAGTT | qPCR; fwd |
| | Pa20117-QPR4 | CCAGAGCAGAGCTTAACAGCA | qPCR; rev |
| *PaMFS5* | Pa22183-CL1 | TGTGGATTTAACATTACCTTACTCCTT | Cloning in pCR4-TOPO vector for sequencing; fwd |
| | Pa22183-CL2 | GAACGCAATTCATATTTTCTAGCTT | Cloning in pCR4-TOPO vector for sequencing; rev |
| | Pa22183-EX1 | TGTCGACATGGCCAACCGCAAAAATG | Expression of gene without stop codon in pIEx-4 vector; fwd |

| | | | |
|---|---|---|---|
| | Pa22183-EX2 | TGCGGCCGCATAATTTGTTAACATTTCTTGTATTTCATG | Expression of gene without stop codon in pIEx-4 vector; rev |
| | Pa22183-S1 | GATGCCAAATTTCGCTGGTAT | Internal sequencing |
| | Pa22183-QPF1 | GGTCGTTTCTGCTGCCTTAG | qPCR; fwd |
| | Pa22183-QPR1 | TTGTGTTCCTCGCACAATTC | qPCR; rev |
| *PaMFS6* | Pa6623-3-CL1 | TGAGTTGCTTTTATCCTGTGCTAT | Cloning in pCR4-TOPO vector for sequencing; fwd |
| | Pa6623-3-CL2 | GATAAGGCTTACAAGCAGTAGATAGGT | Cloning in pCR4-TOPO vector for sequencing; rev |
| | Pa6623-3-EX1 | TGTCGACATGACAGAAAATCGTGTAATTTCG | Expression of gene without stop codon in pIEx-4 vector; fwd |
| | Pa6623-3-EX2 | TGCGGCCGCATAATCGGCTAATATTTGCTGGATT | Expression of gene without stop codon in pIEx-4 vector; rev |
| | Pa6623-3-S1 | CATTTGCATATCGATGGCATT | Internal sequencing |
| | Pa6623-3-QPF1 | TTGGTTGGCTCAGTGTTCAG | qPCR; fwd |
| | Pa6623-3-QPR1 | TCTCCGGTGCTAAATGGAAC | qPCR; rev |
| *PaMFS7* | PaMFS13-CL1 | GATATCAAGCAGTGTATTAAGCACAAT | Cloning in pCR4-TOPO vector for sequencing; fwd |
| | PaMFS13-CL2 | TTTGTTTGAATCCAGTTGACCAA | Cloning in pCR4-TOPO vector for sequencing; rev |
| | PaMFS13-SEQ1 | TCCGGTCTTCAGCATCAAATACT | Internal sequencing |
| | PaMFS13-SEQ2 | CCATAACCAGCGAGCTCTTC | Internal sequencing |
| | PaMFS13-IEX4-1 | ACGCGTCGACATGGAAGATCCCAACGTTTCA | Expression of gene without stop codon in pIEx-4 vector; fwd |
| | PaMFS13-IEX4-2 | ATAAGAATGCGGCCGCAAACTTTTTCAACATCTGCTGTATTT | Expression of gene without stop codon in pIEx-4 vector; rev |
| | PaMFS13-QPF2 | GGGCATTTGTTCGCGTATCT | qPCR; fwd |
| | PaMFS13-QPR2 | GGCACGATGAAGAAGAGCAG | qPCR; rev |
| *PaMFS8* | PaMFS44_5R_a | TATCGCGCCGGCCACACAATTTACGCTA | 5' RACE |
| | PaMFS44_5R_b | TCGACTACGAAGGATGCGACTGTGAAGA | 5' RACE |
| | PaMFS44-CL8 | GTGAATATGTGGGGAAATGGTATATT | Cloning in pCR4-TOPO vector for sequencing; fwd |
| | PaMFS44-CL7 | GGTTCTAATGATTTTGTACAATTTACATAGGTAT | Cloning in pCR4-TOPO vector for sequencing; rev |
| | PaMFS44-IEX4-1 | ACGCGTCGACATGTCCGTCGAAATAGCTGTTA | Expression of gene without stop codon in pIEx-4 vector; fwd |
| | PaMFS44-IEX4-2 | ATAAGAATGCGGCCGCTCTTTCAAGCATTTTCTGTATTTCATC | Expression of gene without stop codon in pIEx-4 vector; rev |
| | PaMFS44-QPF1 | CACTGTCGCATCCTTCGTAGT | qPCR; fwd |

| | | | |
|---|---|---|---|
| | PaMFS44-QPR1 | GCCATTGGAGTTGTTGCAGAG | qPCR; rev, internal sequencing |
| *PaMFS9_ps* | PaMFS50_3R | CCACTGCTAACTAAATACTGTTCCGGTGTTACC | 3' RACE |
| | PaMFS50_5R_a | AGAAACAGAGGTATAGCGGTACTAGTGGTAGA | 5' RACE |
| | PaMFS50_5R_b | GCTACAATGTCACCTGATAAGCTAGTACCTG | 5 'RACE |
| | PaMFS50-CL1 | TATAAGGTGCTGAGTACTAACGGT | Cloning in pCR4-TOPO vector for sequencing; fwd |
| | PaMFS50-CL2 | GGTATTAAGGGGTATTATAAAATGGTATTTG | Cloning in pCR4-TOPO vector for sequencing; rev |
| | PaMFS50_SEQ1 | TGATATATTAGGCAGGAAGCGGT | Internal sequencing |
| *eIF4A* | qPaEiF4a_F | CACGGTGACATGGAGCAAAG | qPCR; fwd |
| | qPaEiF4a_R | ACCTCTGGCCAACAAATCGG | qPCR; rev |
| *RPL13a* | qPaRPL13a_F | CGTTCGTACGTTGGAGAGCA | qPCR; fwd |
| | qPaRPL13a_R | GCTTGGCAACCTTTTCAGTC | qPCR; rev |
| *RPS4e* | qPaRPS4e_F | CGTATTACTGCTGAAGAAGC | qPCR; fwd |
| | qPaRPS4e_R | ATCGTGGGTCACCAAGAACG | qPCR; rev |
| *RPL7* | qPaRPL7_F | AGGCTGAAGGAACGAGATGA | qPCR; fwd |
| | qPaRPL7_R | CTTCGGCTGGAACGTAGAAG | qPCR; rev |
| *IMPI* | T7-IMPI-F2 | TAATACGACTCACTATAGGGAGAGTAATGACAAGTGCTACTGTGAAGAT | Amplification of DNA templates for dsRNA synthesis; fwd |
| | T7-IMPI-R2 | TAATACGACTCACTATAGGGAGAGGGGAGTCAATGCAGGAAAACT | Amplification of DNA templates for dsRNA synthesis; rev (223-bp amplified fragment of *IMPI* using the forward and reverse primers) |

**Supplementary Table 6.** Gene expression variability across tissues among four tested reference genes.

| Target gene | Tissue | Mean Cq value of four biological replicates | Standard deviation of the mean Cq values |
|---|---|---|---|
| | Foregut | 21.70 | |
| | Midgut | 21.96 | |
| *eIF4A* | Hindgut | 21.62 | 0.30 |
| | Malpighian tubules | 21.39 | |
| | Other tissues | 21.17 | |
| | Foregut | 20.81 | |
| | Midgut | 20.44 | |
| *RPL13a* | Hindgut | 20.98 | 0.31 |
| | Malpighian tubules | 20.22 | |
| | Other tissues | 20.49 | |
| | Foregut | 21.42 | |
| | Midgut | 20.39 | |
| *RPL7* | Hindgut | 21.12 | 0.42 |
| | Malpighian tubules | 20.63 | |
| | Other tissues | 20.62 | |
| | Foregut | 21.37 | |
| | Midgut | 20.44 | |
| *RPS4e* | Hindgut | 21.39 | 0.46 |
| | Malpighian tubules | 20.77 | |
| | Other tissues | 20.52 | |

**Supplementary Table 7.** Multiple reaction monitoring (MRM) transitions for compounds determined by LC-MS/MS.

| **Compound** | **Q1 [m/z]** | **Q3 [m/z]** | **CE [eV]** | **Use** |
|---|---|---|---|---|
| 2Prop GLS | 358 | 95.9 | -60 | All samples except those of the pH-dependency experiment using the *Xenopus* oocyte expression system |
| 4MSOB GLS | 435.9 | 95.8 | -60 | |
| 4MTB GLS | 419.9 | 95.9 | -58 | |
| 2PE GLS | 421.81 | 95.9 | -50 | |
| Benzyl GLS | 408 | 95.9 | -60 | |
| 4OHB GLS | 424 | 95.9 | -60 | |
| I3M GLS | 447 | 95.8 | -50 | |
| 3-Butenyl | 372 | 95.9 | -60 | |
| Salicin | 285 | 123 | -18 | |
| Dhurrin | 310 | 179 | -10 | |
| Linamarin (formiate adduct) | 292 | 45 | -26 | |
| Aucubin (formiate adduct) | 391 | 183 | -18 | |
| Catalpol (formiate adduct) | 407 | 199 | -18 | |
| 2Prop GLS | 358 | 97.0[Q] | 22 | Samples of the pH-dependency experiment using the *Xenopus* oocyte expression system |
| | 358 | 75 | 30 | |
| | 358 | 259 | 20 | |
| I3M GLS | 447 | 97.0[Q] | 10 | |
| | 447 | 259 | 10 | |
| | 447 | 205 | 10 | |

[Q]quantifier ion, additional transitions are used for identification only.

**Supplementary Table 8.** *In silico* off-target prediction of the dsRNA designs against the local *P. armoraciae* transcriptome database.

| Target gene | Hit | 1-mismatch count | 2-mismatch count | Sequence annotation |
|---|---|---|---|---|
| *IMPI* | *Parm_de_novo_new_c119* | 0 | 1 | Cytochrome c oxidase assembly protein COX15 homolog |
| | *Parm_de_novo_new_c2603* | 0 | 1 | Serotonin receptor |
| | *Parm_de_novo_new_c5811* | 0 | 1 | Acyl-CoA synthetase family member 4-like |
| | *Parm_de_novo_new_c12340* | 0 | 1 | Zinc finger protein 333 isoform X3 |
| | *Parm_de_novo_new_c14305* | 0 | 1 | Retrovirus-related Pol polyprotein from transposon 412 |
| *PaGTR1* | *Parm_de_novo_new_c331* | 0 | 1 | Nucleolar complex protein 3 homolog |
| | *Parm_de_novo_new_c506* | 0 | 1 | Methyl-CpG-binding domain protein 5 |
| | *Parm_de_novo_new_c521* | 0 | 5 | Mannosyl-oligosaccharide alpha-1,2-mannosidase isoform A |
| | *Parm_de_novo_new_c800* | 0 | 3 | Cyclin-Y |
| | *Parm_de_novo_new_c1397* | 0 | 5 | Actin-related protein 2/3 complex subunit 4 |
| | *Parm_de_novo_new_c1628* | 0 | 1 | Fanconi anemia group M protein |
| | *Parm_de_novo_new_c2850* | 0 | 1 | Sorting nexin-27 |
| | *Parm_de_novo_new_c3233* | 0 | 2 | 3-hydroxy-3-methylglutaryl-coenzyme A reductase |
| | *Parm_de_novo_new_c4725* | 0 | 3 | Max-binding protein MNT-like isoform X1 |
| | *Parm_de_novo_new_c7102* | 0 | 1 | Integrator complex subunit 12 |
| | *Parm_de_novo_new_c7238* | 0 | 1 | Not annotated |
| | *Parm_de_novo_new_c8110* | 0 | 3 | Protein tramtrack, beta isoform-like isoform X1 |
| | *Parm_de_novo_new_c8310* | 0 | 2 | Ribonuclease P protein subunit p29-like |
| | *Parm_de_novo_new_c8643* | 2 | 5 | Arfaptin-2 |
| | *Parm_de_novo_new_c10690* | 0 | 5 | Not annotated |
| | *Parm_de_novo_new_c10867* | 0 | 2 | Not annotated |
| | *Parm_de_novo_new_c14225* | 0 | 6 | Not annotated |
| | *Parm_de_novo_new_c15034* | 0 | 1 | Phosphatidylinositol 4-kinase type 2-beta |
| | *Parm_de_novo_new_c15387* | 0 | 2 | Not annotated |

| | | | | |
|---|---|---|---|---|
| | *Parm_de_novo_new_c17921* | 0 | 1 | Formin 1,2/cappuccino |
| | *Parm_de_novo_new_c23731* | 0 | 1 | Putative gag-pol protein |
| | *Parm_de_novo_new_c23755* | 0 | 1 | Not annotated |
| | *Parm_de_novo_new_c35063* | 0 | 2 | Hypothetical protein GGTG_04705 |
| *PaGTR2* | *Parm_de_novo_new_c1115* | 0 | 3 | Probable ATP-dependent RNA helicase DDX17-like |
| | *Parm_de_novo_new_c2182* | 1 | 2 | Digestive organ expansion factor homolog |
| | *Parm_de_novo_new_c3973* | 0 | 3 | Cullin-3 isoform X1 |
| | *Parm_de_novo_new_c4715* | 0 | 3 | Protein virilizer |
| | *Parm_de_novo_new_c6467* | 0 | 1 | Alpha-tocopherol transfer protein-like isoform |
| | *Parm_de_novo_new_c7416* | 0 | 1 | Not annotated |
| | *Parm_de_novo_new_c16931* | 0 | 1 | Antennal esterase CXE13 |
| | *Parm_de_novo_new_c21176* | 0 | 2 | Not annotated |
| | *Parm_de_novo_new_c23215* | 0 | 1 | Not annotated |
| | *Parm_de_novo_new_c24939* | 0 | 2 | Not annotated |
| | *Parm_de_novo_new_c25100* | 0 | 1 | Not annotated |
| | *Parm_de_novo_new_c32959* | 0 | 4 | Not annotated |
| *PaGTR3* | *Parm_de_novo_new_c441* | 0 | 1 | Titin-like isoform X2 |
| | *Parm_de_novo_new_c1351* | 0 | 2 | Protein lap4-like |
| | *Parm_de_novo_new_c2147* | 0 | 1 | 116 kDa U5 small nuclear ribonucleoprotein component-like |
| | *Parm_de_novo_new_c5806* | 0 | 1 | Serpin peptidase inhibitor 19 |
| | *Parm_de_novo_new_c13822* | 0 | 1 | Slit homolog 3 protein-like |
| | *Parm_de_novo_new_c16099* | 0 | 1 | Encapsulation-relating protein |
| | *Parm_de_novo_new_c21949* | 0 | 1 | 60 kDa lysophospholipase isoform X2 |
| | *Parm_de_novo_new_c22379* | 0 | 1 | Not annotated |
| | *Parm_de_novo_new_c29293* | 0 | 1 | Surface antigen protein |
| *PaGTR5* | *Parm_de_novo_new_c1898* | 0 | 6 | Golgi to ER traffic protein 4 homolog |
| | *Parm_de_novo_new_c19031* | 2 | 2 | Not annotated |
| | *Parm_de_novo_new_c20312* | 0 | 4 | Not annotated |

| | | | | |
|---|---|---|---|---|
| | *Parm_de_novo_new_c8428* | 0 | 3 | Not annotated |
| | *Parm_de_novo_new_c27983* | 1 | 2 | Not annotated |
| | *Parm_de_novo_new_c33950* | 0 | 3 | Solute carrier family 26 member 6-like |
| | *Parm_de_novo_new_c1413* | 0 | 2 | WD repeat and FYVE domain-containing protein 3 |
| | *Parm_de_novo_new_c5502* | 0 | 2 | Not annotated |
| | *Parm_de_novo_new_c10910* | 0 | 2 | Haemolymph juvenile hormone binding protein |
| | *Parm_de_novo_new_c14862* | 0 | 2 | Serine-rich protein |
| | *Parm_de_novo_new_c1217* | 0 | 1 | Vacuolar protein sorting-associated protein 4B |
| | *Parm_de_novo_new_c2400* | 0 | 1 | Forkhead box protein P1-like isoform X9 |
| | *Parm_de_novo_new_c2888* | 0 | 1 | [Pyruvate dehydrogenase (acetyl-transferring)] kinase, mitochondrial |
| | *Parm_de_novo_new_c3461* | 0 | 1 | Midasin-like |
| | *Parm_de_novo_new_c4615* | 0 | 1 | Not annotated |
| | *Parm_de_novo_new_c5804* | 0 | 1 | Small conductance calcium-activated potassium channel protein |
| | *Parm_de_novo_new_c6089* | 0 | 1 | GTP-binding protein Rit1 |
| | *Parm_de_novo_new_c7402* | 0 | 1 | 4-coumarate--CoA ligase 1 |
| | *Parm_de_novo_new_c8215* | 0 | 1 | Not annotated |
| | *Parm_de_novo_new_c8314* | 0 | 1 | Dolichyldiphosphatase 1-like |
| | *Parm_de_novo_new_c9563* | 0 | 1 | Not annotated |
| | *Parm_de_novo_new_c10643* | 0 | 1 | Glycoside hydrolase family 28 |
| | *Parm_de_novo_new_c10861* | 0 | 1 | Not annotated |
| | *Parm_de_novo_new_c11146* | 0 | 1 | Not annotated |
| | *Parm_de_novo_new_c14215* | 0 | 1 | Not annotated |
| | *Parm_de_novo_new_c33204* | 0 | 1 | Major antigen-like |
| | *Parm_de_novo_new_c33479* | 0 | 1 | Not annotated |
| *PaGTR6* | *Parm_de_novo_new_c30521* | 0 | 3 | Cellulose synthase A catalytic subunit 2 [UDP-forming] |
| | *Parm_de_novo_new_c665* | 0 | 1 | Activating signal cointegrator 1 |
| | *Parm_de_novo_new_c5949* | 0 | 1 | Zinc finger and BTB domain-containing protein 49-like |
| | *Parm_de_novo_new_c6478* | 0 | 1 | Ras GTPase-activating protein-binding protein 2 |

| | | | | |
|---|---|---|---|---|
| | *Parm_de_novo_new_c14862* | 0 | 1 | Serine-rich protein |
| *PaGTR7* | *Parm_de_novo_new_c4682* | 0 | 2 | DNA topoisomerase 2-like protein |
| | *Parm_de_novo_new_c330* | 0 | 1 | Not annotated |
| | *Parm_de_novo_new_c3931* | 0 | 1 | Structure-specific endonuclease subunit slx1 |
| | *Parm_de_novo_new_c13139* | 0 | 1 | CDP-diacylglycerol-glycerol-3-phosphate 3-phosphatidyltransferase |
| *PaGTR8* | *Parm_de_novo_new_c2804* | 0 | 2 | Vermiform, isoform G |
| | *Parm_de_novo_new_c24044* | 0 | 2 | RNA polymerase beta subunit-2 |
| | *Parm_de_novo_new_c2121* | 0 | 1 | Sedoheptulokinase |
| | *Parm_de_novo_new_c3301* | 0 | 1 | Serine/threonine-protein kinase polo |
| | *Parm_de_novo_new_c17674* | 0 | 1 | Not annotated |
| | *Parm_de_novo_new_c20798* | 0 | 1 | Not annotated |
| | *Parm_de_novo_new_c25657* | 0 | 1 | Not annotated |

**Supplementary Table 9.** Statistical data.

| **Sample** | | **Transformation** | **Method** | **Statistics** | ***P* value** | **Software** | **Figure** |
|---|---|---|---|---|---|---|---|
| *Pa*GTR1 | 2Prop GLS | Square-root | Two-tailed Student's *t*-test | *t* = -31.068 | < 0.001 | SigmaPlot 14.0 | Fig. 1b |
| | 4MTB GLS | - | Two-tailed Student's *t*-test | *t* = -18.529 | < 0.001 | SigmaPlot 14.0 | |
| | 4OHB GLS | - | Two-tailed Student's *t*-test | *t* = -37.191 | < 0.001 | SigmaPlot 14.0 | |
| | Benzyl GLS | - | Two-tailed Student's *t*-test | *t* = -30.432 | < 0.001 | SigmaPlot 14.0 | |
| | 2PE GLS | - | Two-tailed Student's *t*-test | *t* = -33.231 | < 0.001 | SigmaPlot 14.0 | |
| | I3M GLS | - | Two-tailed Student's *t*-test | *t* = -37.959 | < 0.001 | SigmaPlot 14.0 | |
| *Pa*GTR2 | 2Prop GLS | - | Two-tailed Student's *t*-test | *t* = -21.127 | < 0.001 | SigmaPlot 14.0 | |
| | 4MSOB GLS | - | Two-tailed Student's *t*-test | *t* = -30.704 | < 0.001 | SigmaPlot 14.0 | |
| | 4MTB GLS | - | Two-tailed Student's *t*-test | *t* = -28.404 | < 0.001 | SigmaPlot 14.0 | |
| | 4OHB GLS | - | Two-tailed Student's *t*-test | *t* = -20.774 | < 0.001 | SigmaPlot 14.0 | |
| | Benzyl GLS | - | Two-tailed Student's *t*-test | *t* = -13.320 | < 0.001 | SigmaPlot 14.0 | |
| | 2PE GLS | - | Two-tailed Student's *t*-test | *t* = -16.935 | < 0.001 | SigmaPlot 14.0 | |
| | I3M GLS | - | Two-tailed Student's *t*-test | *t* = -25.150 | < 0.001 | SigmaPlot 14.0 | |
| *Pa*GTR3 | 2Prop GLS | - | Two-tailed Student's *t*-test | *t* = -22.769 | < 0.001 | SigmaPlot 14.0 | |
| | 4MSOB GLS | - | Two-tailed Student's *t*-test | *t* = -15.055 | < 0.001 | SigmaPlot 14.0 | |
| | 4MTB GLS | - | Two-tailed Student's *t*-test | *t* = -64.586 | < 0.001 | SigmaPlot 14.0 | |
| | 4OHB GLS | - | Two-tailed Student's *t*-test | *t* = -43.066 | < 0.001 | SigmaPlot 14.0 | |
| | Benzyl GLS | - | Two-tailed Student's *t*-test | *t* = -61.129 | < 0.001 | SigmaPlot 14.0 | |
| | 2PE GLS | - | Two-tailed Student's *t*-test | *t* = -24.741 | < 0.001 | SigmaPlot 14.0 | |
| | I3M GLS | - | Two-tailed Student's *t*-test | *t* = -20.555 | < 0.001 | SigmaPlot 14.0 | |
| *Pa*GTR4 | 2Prop GLS | - | Two-tailed Student's *t*-test | *t* = -34.895 | < 0.001 | SigmaPlot 14.0 | |
| *Pa*GTR5 | 2Prop GLS | - | Two-tailed Student's *t*-test | *t* = -31.661 | < 0.001 | SigmaPlot 14.0 | |
| | 4MSOB | - | Two-tailed Student's *t*-test | *t* = -10.966 | < 0.001 | SigmaPlot 14.0 | |

| | | | | | | |
|---|---|---|---|---|---|---|
| | GLS | | | | | |
| | 4MTB GLS | - | Two-tailed Student's *t*-test | $t$ = -98.236 | < 0.001 | SigmaPlot 14.0 |
| | Benzyl GLS | - | Two-tailed Student's *t*-test | $t$ = -10.207 | < 0.001 | SigmaPlot 14.0 |
| | 2PE GLS | - | Two-tailed Student's *t*-test | $t$ = -46.917 | < 0.001 | SigmaPlot 14.0 |
| *Pa*GTR6 | 2Prop GLS | - | Two-tailed Student's *t*-test | $t$ = -30.115 | < 0.001 | SigmaPlot 14.0 |
| | 4MTB GLS | - | Generalized least squares | $LR$ = 13.019 | < 0.001 | R 3.5.1 |
| | Benzyl GLS | - | Two-tailed Student's *t*-test | $t$ = -33.478 | < 0.001 | SigmaPlot 14.0 |
| | 2PE GLS | Square-root | Two-tailed Student's *t*-test | $t$ = -56.933 | < 0.001 | SigmaPlot 14.0 |
| *Pa*GTR7 | 2Prop GLS | Square-root | Two-tailed Student's *t*-test | $t$ = -127.779 | < 0.001 | SigmaPlot 14.0 |
| *Pa*GTR8 | 4MTB GLS | - | Two-tailed Student's *t*-test | $t$ = -49.878 | < 0.001 | SigmaPlot 14.0 |
| | 4OHB GLS | - | Two-tailed Student's *t*-test | $t$ = -50.592 | < 0.001 | SigmaPlot 14.0 |
| | Benzyl GLS | - | Two-tailed Student's *t*-test | $t$ = -42.186 | < 0.001 | SigmaPlot 14.0 |
| | 2PE GLS | - | Two-tailed Student's *t*-test | $t$ = -61.485 | < 0.001 | SigmaPlot 14.0 |
| *Pa*GTR9 | 2Prop GLS | - | Two-tailed Student's *t*-test | $t$ = -84.908 | < 0.001 | SigmaPlot 14.0 |
| | 4MTB GLS | - | Two-tailed Student's *t*-test | $t$ = -66.857 | < 0.001 | SigmaPlot 14.0 |
| | Benzyl GLS | - | Two-tailed Student's *t*-test | $t$ = -134.168 | < 0.001 | SigmaPlot 14.0 |
| | 2PE GLS | - | Two-tailed Student's *t*-test | $t$ = -99.949 | < 0.001 | SigmaPlot 14.0 |
| | I3M GLS | $Log_{10}$ | Two-tailed Student's *t*-test | $t$ = -26.343 | < 0.001 | SigmaPlot 14.0 |
| *Pa*GTR10 | 2Prop GLS | Square-root | Two-tailed Student's *t*-test | $t$ = -61.124 | < 0.001 | SigmaPlot 14.0 |
| | 4MTB GLS | - | Two-tailed Student's *t*-test | $t$ = -74.811 | < 0.001 | SigmaPlot 14.0 |
| *Pa*GTR11 | 2Prop GLS | - | Two-tailed Student's *t*-test | $t$ = -51.331 | < 0.001 | SigmaPlot 14.0 |
| | 4MSOB GLS | - | Two-tailed Student's *t*-test | $t$ = -49.142 | < 0.001 | SigmaPlot 14.0 |
| | 4MTB GLS | - | Two-tailed Student's *t*-test | $t$ = -44.957 | < 0.001 | SigmaPlot 14.0 |
| | 4OHB GLS | - | Two-tailed Student's *t*-test | $t$ = -42.624 | < 0.001 | SigmaPlot 14.0 |
| | Benzyl GLS | - | Two-tailed Student's *t*-test | $t$ = -57.609 | < 0.001 | SigmaPlot 14.0 |

| | | | | | | | |
|---|---|---|---|---|---|---|---|
| | 2PE GLS | - | Two-tailed Student's *t*-test | *t* = -66.217 | < 0.001 | SigmaPlot 14.0 | |
| *Pa*GTR12 | 2Prop GLS | - | Two-tailed Student's *t*-test | *t* = -1.118 | 0.326 | SigmaPlot 14.0 | |
| | 4MSOB GLS | Square-root | Two-tailed Student's *t*-test | *t* = 3.092 | 0.037 | SigmaPlot 14.0 | |
| | 4MTB GLS | - | Two-tailed Student's *t*-test | *t* = -14.246 | < 0.001 | SigmaPlot 14.0 | |
| | Benzyl GLS | - | Two-tailed Student's *t*-test | *t* = -1.138 | 0.319 | SigmaPlot 14.0 | |
| | 2PE GLS | - | Two-tailed Student's *t*-test | *t* = -3.775 | 0.02 | SigmaPlot 14.0 | |
| *Pa*GTR13 | 4MSOB GLS | - | Two-tailed Student's *t*-test | *t* = -34.924 | < 0.001 | SigmaPlot 14.0 | |
| | 4MTB GLS | - | Two-tailed Student's *t*-test | *t* = -64.164 | < 0.001 | SigmaPlot 14.0 | |
| | 4OHB GLS | - | Two-tailed Student's *t*-test | *t* = -22.168 | < 0.001 | SigmaPlot 14.0 | |
| | 2PE GLS | - | Two-tailed Student's *t*-test | *t* = -29.928 | < 0.001 | SigmaPlot 14.0 | |
| *Pa*MFS2 | Aucubin | - | Two-tailed Student's *t*-test | *t* = -26.754 | < 0.001 | SigmaPlot 14.0 | |
| *Pa*MFS6 | Catalpol | Square-root | Two-tailed Student's *t*-test | *t* = -15.421 | < 0.001 | SigmaPlot 14.0 | |
| | Aucubin | - | Two-tailed Student's *t*-test | *t* = -18.699 | < 0.001 | SigmaPlot 14.0 | |
| Gene expression level | | - | Mann-Whitney *U* test | *U* = 0.000 | 0.008 | SigmaPlot 14.0 | Fig. 2a |
| I3M GLS | | - | Two-tailed Student's *t*-test | *t* = 10.975 | < 0.001 | SigmaPlot 14.0 | Fig. 2b |
| 1MOI3M GLS | | - | Mann-Whitney *U* test | *U* = 0.000 | < 0.001 | | |
| I3M GLS | Day 1 | Square-root | Two-tailed Student's *t*-test | *t* = -4.947 | < 0.001 | SigmaPlot 14.0 | Fig. 2c |
| | Day 2 | - | Generalized least squares | *LR* = 1.689 | 0.194 | R 3.5.1 | |
| | Day 3 | - | Two-tailed Student's *t*-test | *t* = -1.585 | 0.13 | SigmaPlot 14.0 | |
| | Day 4 | - | Two-tailed Student's *t*-test | *t* = -0.258 | 0.799 | SigmaPlot 14.0 | |
| | Day 5 | - | Two-tailed Student's *t*-test | *t* = 0.365 | 0.72 | SigmaPlot 14.0 | |
| 1MOI3M GLS | Day 1[1] | - | - | - | - | - | |
| | Day 2 | Square-root | Two-tailed Student's *t*-test | *t* = -0.483 | 0.635 | SigmaPlot 14.0 | |
| | Day 3 | - | Two-tailed Student's *t*-test | *t* = -3.204 | 0.005 | SigmaPlot 14.0 | |
| | Day 4 | - | Two-tailed Student's *t*-test | *t* = -1.970 | 0.066 | SigmaPlot 14.0 | |
| | Day 5 | - | Two-tailed Student's *t*-test | *t* = 0.651 | 0.524 | SigmaPlot 14.0 | |

| | | | | | | |
|---|---|---|---|---|---|---|
| 2Prop GLS | $Log_{10}$ | Two-tailed Student's *t*-test | *t* = -28.769 | < 0.001 | SigmaPlot 14.0 | Fig. 2d |
| 4MSOB GLS | Square-root | Two-tailed Student's *t*-test | *t* = -3.630 | 0.011 | | |
| 4MTB GLS | $Log_{10}$ | Two-tailed Student's *t*-test | *t* = -22.118 | < 0.001 | | |
| 4OHB GLS | $Log_{10}$ | Two-tailed Student's *t*-test | *t* = -71.513 | < 0.001 | | |
| Benzyl GLS | $Log_{10}$ | Two-tailed Student's *t*-test | *t* = -66.466 | < 0.001 | | |
| 2PE GLS | Square-root | Two-tailed Student's *t*-test | *t* = -47.387 | < 0.001 | | |
| I3M GLS | Square-root | Two-tailed Student's *t*-test | *t* = -55.369 | < 0.001 | | |
| Transport activity | - | One-way ANOVA | *F*= 30.803 | < 0.001 | SigmaPlot 14.0 | Fig. 2e |
| Transport activity | Square-root | One-way ANOVA | *F*= 39.664 | < 0.001 | SigmaPlot 14.0 | Fig. 2f |
| Glucosides in beetles 30 min after the injection vs. after feeding for one day: 2Prop GLS | - | Two-tailed Student's *t*-test | *t* = 1.354 | 0.192 | SigmaPlot 14.0 | - |
| 4OHB GLS | - | Two-tailed Student's *t*-test | *t* = 0.929 | 0.365 | | |
| Salicin | Square-root | Two-tailed Student's *t*-test | *t* = 7.307 | < 0.001 | | |
| Linamarin | - | Mann-Whitney *U* test | *U* = 0.000 | < 0.001 | | |
| Catalpol | Square-root | Two-tailed Student's *t*-test | *t* = 3.544 | 0.002 | | |
| Glucosides in beetles | - | Generalized least squares | *LR* = 117.719 | < 0.001 | R 3.5.1 | Fig. 3a |
| Glucosides in feces | - | Generalized least squares | *LR* = 52.4334 | < 0.001 | | |
| Bathing saline vs. Malpighian tubules: 2Prop GLS | - | Two-tailed Student's *t*-test | *t* = 4.577 | 0.001 | SigmaPlot 14.0 | Fig. 3b |
| 4MSOB GLS | - | | *t* = 4.189 | 0.002 | | |
| 4OHB GLS | - | | *t* = 3.865 | 0.003 | | |
| 2PE GLS | - | | *t* = 5.858 | < 0.001 | | |
| I3M GLS | - | | *t* = 3.446 | 0.006 | | |
| Salicin | - | | *t* = -3.765 | 0.004 | | |
| Linamarin | - | | *t* = -4.168 | 0.002 | | |
| Catalpol | - | | *t* = -4.888 | < 0.001 | | |
| Malpighian tubules vs. excretion: 2Prop GLS | - | | *t* = 5.492 | < 0.001 | | |
| 4MSOB GLS | - | | *t* = 3.250 | 0.009 | | |

| | | | | | | | |
|---|---|---|---|---|---|---|---|
| fluid | 4OHB GLS | - | | *t* = 3.741 | 0.004 | | |
| | 2PE GLS | - | | *t* = 6.026 | < 0.001 | | |
| | I3M GLS | - | | *t* = 0.585 | 0.571 | | |
| | Salicin | - | | *t* = -11.754 | < 0.001 | | |
| | Linamarin | - | | *t* = -0.746 | 0.473 | | |
| | Catalpol | - | | *t* = 3.543 | 0.005 | | |
| *PaGTR5* | | $Log_{10}$ | Two-tailed Student's *t*-test | *t* = 8.711 | < 0.001 | SigmaPlot 14.0 | Fig. 4a |
| *PaGTR6* | | - | Two-tailed Student's *t*-test | *t* = 4.371 | 0.001 | SigmaPlot 14.0 | |
| *PaGTR7* | | Square-root | Two-tailed Student's *t*-test | *t* = 9.197 | < 0.001 | SigmaPlot 14.0 | |
| *PaGTR8* | | - | Mann-Whitney *U* test | *U* = 0.000 | 0.002 | SigmaPlot 14.0 | |
| 2Prop GLS | | - | Two-tailed Student's *t*-test | *t* = 2.825 | 0.011 | SigmaPlot 14.0 | Fig. 4b |
| 2Prop GLS | Day 2 | - | Mann-Whitney *U* test | *U* = 39.000 | 0.427 | SigmaPlot 14.0 | Fig. 4c |
| | Day 3 | Square-root | Two-tailed Student's *t*-test | *t* = -1.656 | 0.115 | SigmaPlot 14.0 | |
| | Day 4 | Square-root | Two-tailed Student's *t*-test | *t* = -2.113 | 0.049 | SigmaPlot 14.0 | |
| | Day 5 | - | Two-tailed Student's *t*-test | *t* = -2.304 | 0.033 | SigmaPlot 14.0 | |
| | Day 6 | - | Two-tailed Student's *t*-test | *t* = -2.933 | 0.009 | SigmaPlot 14.0 | |
| *PaGTR2* | | - | Two-tailed Student's *t*-test | *t* = -1.546 | 0.161 | SigmaPlot 14.0 | Supplementary Figure 3a |
| *PaGTR3* | | - | Mann-Whitney *U* test | *U* = 11.000 | 0.841 | | |
| *PaGTR9* | | - | Two-tailed Student's *t*-test | *t* = 0.910 | 0.389 | | |
| *PaGTR10* | | - | Two-tailed Student's *t*-test | *t* = 0.866 | 0.412 | | |
| Day 1 | | - | Two-tailed Student's *t*-test | *t* = -1.907 | 0.073 | SigmaPlot 14.0 | Supplementary Figure 3b |
| Day 2 | | - | Mann-Whitney *U* test | *U* = 49.000 | 0.97 | | |
| Day 3 | | - | Two-tailed Student's *t*-test | *t* = -0.779 | 0.446 | | |
| Day 4 | | - | Two-tailed Student's *t*-test | *t* = 0.114 | 0.91 | | |
| Day 5 | | - | Two-tailed Student's *t*-test | *t* = 1.403 | 0.178 | | |
| 2Prop GLS | Day 1 | - | Two-tailed Student's *t*-test | *t* = -0.293 | 0.773 | SigmaPlot 14.0 | Supplementary Figure 4 |
| | Day 2 | - | Two-tailed Student's *t*-test | *t* = 0.517 | 0.612 | SigmaPlot 14.0 | |
| | Day 3 | - | Two-tailed Student's *t*-test | *t* = 0.187 | 0.854 | SigmaPlot 14.0 | |

| | | | | | | |
|---|---|---|---|---|---|---|
| | Day 4 | Square-root | Two-tailed Student's $t$-test | $t$ = 0.897 | 0.383 | SigmaPlot 14.0 |
| | Day 5 | - | Two-tailed Student's $t$-test | $t$ = 0.650 | 0.524 | SigmaPlot 14.0 |
| 3MSOP GLS | Day 1 | - | Mann-Whitney $U$ test | $U$ = 49.000 | 0.968 | SigmaPlot 14.0 |
| | Day 2 | - | Two-tailed Student's $t$-test | $t$ = -0.231 | 0.82 | SigmaPlot 14.0 |
| | Day 3 | - | Two-tailed Student's $t$-test | $t$ = 0.922 | 0.369 | SigmaPlot 14.0 |
| | Day 4 | Square-root | Two-tailed Student's $t$-test | $t$ = -0.256 | 0.801 | SigmaPlot 14.0 |
| | Day 5 | - | Two-tailed Student's $t$-test | $t$ = -0.500 | 0.623 | SigmaPlot 14.0 |
| 4MSOB GLS | Day 1 | Square-root | Two-tailed Student's $t$-test | $t$ = -0.977 | 0.341 | SigmaPlot 14.0 |
| | Day 2 | - | Two-tailed Student's $t$-test | $t$ = -1.215 | 0.24 | SigmaPlot 14.0 |
| | Day 3 | Square-root | Two-tailed Student's $t$-test | $t$ = 0.0495 | 0.961 | SigmaPlot 14.0 |
| | Day 4 | Square-root | Two-tailed Student's $t$-test | $t$ = 0.276 | 0.786 | SigmaPlot 14.0 |
| | Day 5 | - | Two-tailed Student's $t$-test | $t$ = 0.954 | 0.353 | SigmaPlot 14.0 |
| 5MSOP GLS | Day 1 | - | Two-tailed Student's $t$-test | $t$ = -1.253 | 0.226 | SigmaPlot 14.0 |
| | Day 2 | - | Two-tailed Student's $t$-test | $t$ = -0.461 | 0.65 | SigmaPlot 14.0 |
| | Day 3 | - | Two-tailed Student's $t$-test | $t$ = -0.105 | 0.918 | SigmaPlot 14.0 |
| | Day 4 | - | Two-tailed Student's $t$-test | $t$ = 0.580 | 0.57 | SigmaPlot 14.0 |
| | Day 5 | - | Two-tailed Student's $t$-test | $t$ = 1.112 | 0.281 | SigmaPlot 14.0 |
| 7MSOH GLS | Day 1[1] | - | - | - | - | - |
| | Day 2 | - | Mann-Whitney $U$ test | $U$ = 47.000 | 0.845 | SigmaPlot 14.0 |
| | Day 3 | - | Two-tailed Student's $t$-test | $t$ = -0.176 | 0.863 | SigmaPlot 14.0 |
| | Day 4 | - | Mann-Whitney $U$ test | $U$ = 37.500 | 0.825 | SigmaPlot 14.0 |
| | Day 5 | - | Mann-Whitney $U$ test | $U$ = 44.000 | 0.675 | SigmaPlot 14.0 |
| 8MSOO GLS | Day 1[1] | - | - | - | - | - |
| | Day 2 | - | Mann-Whitney $U$ test | $U$ = 43.000 | 0.564 | SigmaPlot 14.0 |
| | Day 3 | - | Mann-Whitney $U$ test | $U$ = 44.000 | 0.671 | SigmaPlot 14.0 |
| | Day 4 | - | Mann-Whitney $U$ test | $U$ = 34.000 | 0.595 | SigmaPlot 14.0 |
| | Day 5 | - | Mann-Whitney $U$ test | $U$ = 37.500 | 0.358 | SigmaPlot 14.0 |
| 4MOI3M GLS | Day 1 | - | Two-tailed Student's $t$-test | $t$ = -2.550 | 0.02 | SigmaPlot 14.0 |
| | Day 2 | - | Two-tailed Student's $t$-test | $t$ = 1.496 | 0.152 | SigmaPlot 14.0 |

| | | | | | | | |
|---|---|---|---|---|---|---|---|
| | Day 3 | - | Two-tailed Student's *t*-test | $t = -0.217$ | 0.831 | SigmaPlot 14.0 | |
| | Day 4 | - | Two-tailed Student's *t*-test | $t = -0.190$ | 0.851 | SigmaPlot 14.0 | |
| | Day 5 | - | Two-tailed Student's *t*-test | $t = 1.433$ | 0.169 | SigmaPlot 14.0 | |
| *PaGTR9* | | - | Two-tailed Student's *t*-test | $t = 1.403$ | 0.191 | SigmaPlot 14.0 | Supplementary Figure 7a |
| *PaGTR10* | | - | Two-tailed Student's *t*-test | $t = -0.0736$ | 0.943 | | |
| Day 2 | | - | Two-tailed Student's *t*-test | $t = 1.367$ | 0.188 | SigmaPlot 14.0 | Supplementary Figure 7b |
| Day 3 | | Square-root | Two-tailed Student's *t*-test | $t = -1.044$ | 0.310 | | |
| Day 4 | | - | Two-tailed Student's *t*-test | $t = -1.555$ | 0.137 | | |
| Day 5 | | - | Two-tailed Student's *t*-test | $t = 0.181$ | 0.859 | | |
| Day 6 | | - | Two-tailed Student's *t*-test | $t = 0.214$ | 0.833 | | |
| 3MSOP GLS | Day 2[1] | - | - | - | - | - | Supplementary Figure 8 |
| | Day 3 | - | Two-tailed Student's *t*-test | $t = -1.987$ | 0.062 | SigmaPlot 14.0 | |
| | Day 4 | - | Two-tailed Student's *t*-test | $t = -0.675$ | 0.509 | SigmaPlot 14.0 | |
| | Day 5 | - | Two-tailed Student's *t*-test | $t = 0.762$ | 0.456 | SigmaPlot 14.0 | |
| | Day 6 | - | Two-tailed Student's *t*-test | $t = 0.894$ | 0.383 | SigmaPlot 14.0 | |
| 4MSOB GLS | Day 2 | - | Mann-Whitney *U* test | $U = 39.000$ | 0.427 | SigmaPlot 14.0 | |
| | Day 3 | - | Mann-Whitney *U* test | $U = 39.000$ | 0.427 | SigmaPlot 14.0 | |
| | Day 4 | Square-root | Two-tailed Student's *t*-test | $t = -0.261$ | 0.797 | SigmaPlot 14.0 | |
| | Day 5 | Square-root | Two-tailed Student's *t*-test | $t = 0.808$ | 0.430 | SigmaPlot 14.0 | |
| | Day 6 | - | Two-tailed Student's *t*-test | $t = 1.619$ | 0.123 | SigmaPlot 14.0 | |
| 5MSOP GLS | Day 2[1] | - | - | - | - | - | |
| | Day 3[1] | - | - | - | - | - | |
| | Day 4[1] | - | - | - | - | - | |
| | Day 5 | - | Mann-Whitney *U* test | $U = 28.500$ | 0.090 | SigmaPlot 14.0 | |
| | Day 6 | - | Two-tailed Student's *t*-test | $t = 1.505$ | 0.150 | SigmaPlot 14.0 | |
| 7MSOH GLS | Day 2[1] | - | - | - | - | - | |
| | Day 3[1] | - | - | - | - | - | |

| | | | | | | | |
|---|---|---|---|---|---|---|---|
| | Day 4[1] | - | - | - | - | - | |
| | Day 5[1] | - | - | - | - | - | |
| | Day 6[1] | - | - | - | - | - | |
| 8MSOO GLS | Day 2[1] | - | - | - | - | - | |
| | Day 3[1] | - | - | - | - | - | |
| | Day 4[1] | - | - | - | - | - | |
| | Day 5[1] | - | - | - | - | - | |
| | Day 6[1] | - | - | - | - | - | |
| I3M GLS | Day 2[1] | - | - | - | - | - | |
| | Day 3 | - | Mann-Whitney *U* test | *U* = 24.000 | 0.043 | SigmaPlot 14.0 | |
| | Day 4 | - | Mann-Whitney *U* test | *U* = 33.000 | 0.211 | SigmaPlot 14.0 | |
| | Day 5 | - | Two-tailed Student's *t*-test | *t* = 3.170 | 0.005 | SigmaPlot 14.0 | |
| | Day 6 | - | Two-tailed Student's *t*-test | *t* = 2.440 | 0.025 | SigmaPlot 14.0 | |
| 4MOI3M GLS | Day 2 | - | Two-tailed Student's *t*-test | *t* = 2.304 | 0.033 | SigmaPlot 14.0 | |
| | Day 3 | - | Two-tailed Student's *t*-test | *t* = 0.149 | 0.883 | SigmaPlot 14.0 | |
| | Day 4 | - | Two-tailed Student's *t*-test | *t* = -2.835 | 0.011 | SigmaPlot 14.0 | |
| | Day 5 | - | Two-tailed Student's *t*-test | *t* = 3.252 | 0.004 | SigmaPlot 14.0 | |
| | Day 6 | - | Two-tailed Student's *t*-test | *t* = -0.0386 | 0.970 | SigmaPlot 14.0 | |
| *PaGTR2* | | Square-root | Two-tailed Student's *t*-test | *t* = 9.832 | < 0.001 | SigmaPlot 14.0 | Supplementary Figure 9a |
| *PaGTR3* | | $Log_{10}$ | Two-tailed Student's *t*-test | *t* = 5.679 | < 0.001 | | |
| Accumulation of the ingested glucosinolates | | - | Two-tailed Student's *t*-test | *t* = -1.365 | 0.185 | SigmaPlot 14.0 | Supplementary Figure 9b |

[1]Statistical analysis was not conducted if glucosinolate was only detected in one or two replicates.
